# Oncolytic measles virus reprograms the tumor microenvironment in a vascularized mesothelioma-on-chip model

**DOI:** 10.64898/2026.05.12.724508

**Authors:** Nikita Rajkumari, Mégane Willems, Judith Fresquet, Elise Douillard, Magali Devic, Hortense Perdrieau, Delphine Fradin, Jean-François Fonteneau, Nicolas Boisgerault, Isabelle Corre, Lucas Treps, Boudewijn van der Sanden, Christophe Blanquart

**Affiliations:** Nantes Université, Inserm UMR 1307, CNRS UMR 6075, Université d’Angers, CRCI2NA, F-44000 Nantes, France; Platform of Intravital Microscopy, TIMC, CNRS UMR 5525, Université Grenoble Alpes, Grenoble INP, INSERM, Grenoble, France

## Abstract

Pleural mesothelioma (PM) is a rare, aggressive cancer primarily caused by asbestos exposure and remains resistant to conventional chemotherapy. Although dual immune checkpoint inhibition (anti-PD-1/anti-CTLA-4) is now approved as first-line therapy, clinical benefit is limited to a small subset of patients, necessitating the need for alternative strategies. Oncolytic viruses (OVs) represent a promising approach as they selectively infect and lyse tumor cells while reprogramming the immunosuppressive tumor microenvironment (TME) into an immunostimulatory state. In PM, we previously showed that the attenuated Schwarz strain of measles virus (MV) oncolytic activity is mainly dependent on alterations in the type I interferon (IFN-I) pathway, rendering tumor cells sensitive to infection. Recently, we showed that monocytes/macrophages exposed to MV produce IFN-I, which protects PM cells via paracrine IFNAR signaling. This underscores the necessity of modeling the TME to accurately evaluate OV efficacy. Conventional rodent models are non-permissive to MV, and availability of fresh human PM tissue is scarce. We therefore developed a humanized 3D “vascularized mesothelioma-on-chip” (VMOC) model using microfluidic chips. It comprises two perfusable endothelial-lined parental vessels flanking a central secondary microvascular network (MVN), generated using human umbilical vein endothelial cells (HUVECs) embedded in fibrin and co-cultured alongside PM cells and primary human lung fibroblasts (hLFs). We characterized the integrity and functionality of the endothelial compartment as well as the cellular heterogeneity in VMOC using single-cell RNA sequencing. After administration of MV via the endothelial network, we observed infection and death of PM cells in addition to a strong activation of the type I interferon pathway and production of multiple inflammatory mediators. The VMOC model enables *in vitro* study of both MV infection and TME reprogramming, paving the way for a better understanding of the role of the TME in the response to treatment and for supporting the development of more personalized, targeted therapies for PM.

## Introduction

Pleural mesothelioma (PM) is a rare, aggressive malignancy primarily caused by asbestos exposure that arises from the mesothelial cells of the pleura. For the past 20 years, the standard first-line regimen for PM has consisted of platinum plus pemetrexed, with a median overall survival of ∼12 months. In 2020, combined anti-PD-1/anti-CTLA-4 immune checkpoint inhibition was approved as first-line therapy for unresectable PM; however, durable responses remain limited to only a subset of patients (1). Despite extensive preclinical effort, most candidate therapies that demonstrated promising efficacy in preclinical models have failed to translate into meaningful clinical benefit (2,3). These challenges not only underscore the need for alternative therapeutic strategies but also for better preclinical models.

Among the various innovative therapeutic approaches under investigation, oncolytic viruses (OVs) are of particular interest because they can selectively infect and lyse cancer cells while sparing healthy tissue and reprogram the immunosuppressive tumor microenvironment (TME) toward an immunologically active state. The attenuated Schwarz strain of measles virus (MV), an enveloped single-stranded RNA paramyxovirus, has been investigated as an oncolytic agent in PM. MV preferentially uses CD46 for cell entry (4,5), which is frequently overexpressed on PM and other tumor cells to evade complement-mediated lysis (6,7), whereas wild-type strains rely on CD150/SLAM and Nectin 4 (8,9). However, receptor usage alone does not account for oncolytic selectivity. Previous studies from our team have demonstrated that permissiveness of PM cells to MV replication is primarily driven by alterations in the innate antiviral type I interferon (IFN-I) pathway, particularly homozygous deletions of all IFN-I genes (6,10). In addition, infection of PM cells by MV has been shown to improve immunogenicity by inducing dendritic cell maturation, enabling cross-priming of tumor-associated antigen to T cells, and promoting antigen spreading (11–13). In immunocompetent mouse tumor models, MV treatment has been associated with increased intratumoral T-cell infiltration, reduced tumor growth and prolonged survival (14,15).

Clinical evaluations of MV using the Edmonston-Zagreb strain have been performed on multiple cancers, including PM, ovarian cancer, cutaneous T-cell lymphoma, multiple myeloma, and glioblastoma, which have reported acceptable safety profiles (16). However, clinical responses have been heterogeneous so far. This heterogeneity points to context-specific determinants of MV efficacy, including the complexity of the tumor ecosystem. This hypothesis is supported by recent work from our team showing that monocytes/macrophages produce IFN-I in response to MV infection, which reduces viral replication in neighboring virus-sensitive PM cells through paracrine activation of the interferon-α/β receptor (IFNAR) signaling pathway (17). Although many PM cells cannot produce IFN-I themselves, some remain responsive to exogenous IFN-I. Therefore, integrating relevant components from the TME in the evaluation of MV oncolytic activity is essential.

Rodents are not naturally permissive to MV infection, as they lack a functional equivalent of the human CD46 receptor and require genetic engineering, including CD46-transgenic or IFNAR-knockout mice models (18,19), to become permissive. However, these modifications do not fully reproduce the physiological context of human infection, thus limiting their ability to model MV infection. On the other hand, the use of fresh patient tissues is limited due to the rarity of mesothelioma and the small proportion of patients eligible for surgery. In recent years, three-dimensional *in vitro* cancer models have emerged as a promising approach to study the TME, offering an alternative to conventional experimental systems. They provide spatial control over cellular organization and enable physiologically relevant cell-cell and cell-matrix interactions. Models incorporating a functional vascular compartment and other stromal components have already been used to study mechanisms of viral delivery and infection (20–23). Thus, the objective of this study was to develop a TME-relevant vascularized mesothelioma-on-chip (VMOC) model using a three-channel chip, comprising endothelial-lined parental vessels (PV) in the two lateral media channels (LCs) connected to a perfusable microvascular network (MVN) embedded in fibrin gel in the central gel channel (CC). The two endothelial compartments were optimized and characterized independently before generating them in a unified system. The PM cell line Meso13 and primary human lung fibroblasts (hLFs) were incorporated within the fibrin gel to recapitulate key tumor-stromal interactions. Cellular heterogeneity in the model was characterized using single-cell RNA Sequencing (scRNA-seq). Different methods to establish stable MV infection were evaluated, and administration via the PV was selected to mimic intravenous injection. Finally, the consequences of MV infection on tumor cell death as well as on the inflammatory and anti-viral responses were evaluated within the model.

## Material & Methods

### Cell Culture

#### Human umbilical vein endothelial cells (HUVECs)

HUVECs were freshly isolated from umbilical cords obtained from consenting donors at the CHU maternity (Nantes, France) as previously described (24). Informed consent was obtained from all subjects. After collection, they were flushed with 20 mL of phosphate-buffered saline (PBS) to remove residual blood. The bottom of the cord was clamped with a sterile clip. Ten mL of digestion mix containing 0.2% collagenase type I (Life Technologies, cat. no. 17100-17), 2 mM CaCl_2_, 100 IU/mL penicillin, and 0.1 mg/mL streptomycin (Gibco, cat. no. 15140) in 0.9% NaCl was injected from the top of the cord. The cord was incubated for 13 minutes at 37°C. After incubation, the umbilical cord was unclipped, and the digestion mix was collected. The vein was rinsed with M199 medium to collect residual cells. The cell suspension was centrifuged at 300 x g for 5 minutes. The supernatant was discarded, and cells were cultured in M199 medium supplemented with 200 IU/mL penicillin and 0.2 mg/mL streptomycin. T75 tissue culture flasks pre-coated with 0.1% gelatin (Sigma, cat. no. G9391-100G) were used for all HUVEC culture. Cultures were maintained at 37 °C with 5% CO₂. After the first passage, the cells were cultured in EBM2 (PromoCell, cat. no. C-22211), supplemented with the provided supplement mix (PromoCell, cat. no. C-39216), hereafter referred to as supplemented EBM2. The cells were used from passage 2 to 6. For live microscopy experiments, HUVECs were further transduced with lentiviral vectors expressing the red fluorescent protein mRuby2.

#### Human Lung Fibroblast (hLFs)

Healthy lung tissue biopsies were obtained from consenting patients undergoing surgical resection for non-small cell lung carcinoma at Nantes university hospital (France). According to a previously described protocol from our group (24), tissue samples were rinsed with PBS and minced into fragments <1mm³. Samples were digested with 5 mL of digestion medium containing Dulbecco’s Modified Eagle Medium (DMEM) (4.5 g/l glucose) supplemented with 200 IU/mL penicillin, 0.2 mg/mL streptomycin, 0.1% collagenase type I (Life Technologies, cat. no. 17100-17), 0.1 % collagenase type II (Life Technologies, cat. No. 17101-015), and 2.5 U/mL dispase (Life Technologies, cat. no. 17105-041). Digestion was carried out at 37 °C with manual agitation every 5 minutes. After 30 minutes of digestion, 10 mL of cold PBS solution containing 0.1 % bovine serum albumin (BSA) (Clearline, cat. no. 509387) was added. The suspension was then filtered through a 100 µm cell strainer (Biologix, cat. no. 15-1100). Cells were centrifugated at 300 x g for 5 minutes and resuspended in a 1:1 mixture of supplemented EBM2 and DMEM. Cells were cultured in 0.1% gelatin coated culture plates. After 24 hours, non-adherent cells were removed, and the medium was replaced with EBM2. Once cells reached confluence, a CD31 selection (CD31-targeted magnetic cell sorting, MACS) was carried out to remove the endothelial cells and isolate fibroblasts from the negative population. The CD31-negative (fibroblastic) population was characterized by flow cytometry and cultured in DMEM (Gibco) supplemented with 2 mM L-glutamine, 100 IU/mL penicillin, 0.1 mg/mL streptomycin, and 10% heat-inactivated fetal calf serum (Biosera), hereafter referred to as supplemented DMEM. Cell cultures were maintained at 37 °C in a 5% CO₂ humidified incubator. The cells were used from passage 2 to 9.

#### Pleural Mesothelioma cells

The PM cell line, Meso13, was established in our laboratory from the pleural effusion of a patient. It is part of a certified biocollection (Ministère de l’Enseignement Supérieur et de la Recherche, DC-2017-2987; CNIL, 1657097) and has been characterized for its mutational status using targeted sequencing (25). Meso13 cells were transduced with lentiviral vector (pLenti6.1) encoding the blue fluorescent protein 2 (Meso13-BFP) and lentiviral vector (pMX2.1) encoding luciferase (Meso13-luc) for live imaging and viability assays, respectively. Cells were cultured in RPMI-1640 medium (Gibco) supplemented with 2 mM L-glutamine, 100 IU/mL penicillin, 0.1 mg/mL streptomycin, and 10% heat-inactivated fetal calf serum (Biosera). Cell culture was maintained at 37 °C with 5 % CO₂ in a humidified incubator.

### Microfluidic chip

The idenTX microfluidic chip (AIM Biotech, cat. no. DAX-1) and its holder (AIM Biotech, cat. no. HOL-1) were used for all experiments (Figure 1A). The device is manufactured in a standard microscope slide format (approximately 75 × 25 mm) and comprises three independent systems each constituted of three parallel microchannels: a central gel channel (∼1.3 mm wide) flanked by two lateral media channels (∼0.5 mm wide each). All channels are approximately 10.5 mm in length and 0.25 mm in height. The central gel channel is separated from the lateral media channels by arrays of trapezoidal micropillars (0.05 mm wide, ∼0.10 mm spacing) that retain the hydrogel within the central gel channel. Each independent system contains six inlets: four inlets connected to reservoirs feeding the two lateral media channels (two inlets per lateral media channel) and two inlets connected to the central gel channel without reservoirs.

**Figure 1.**
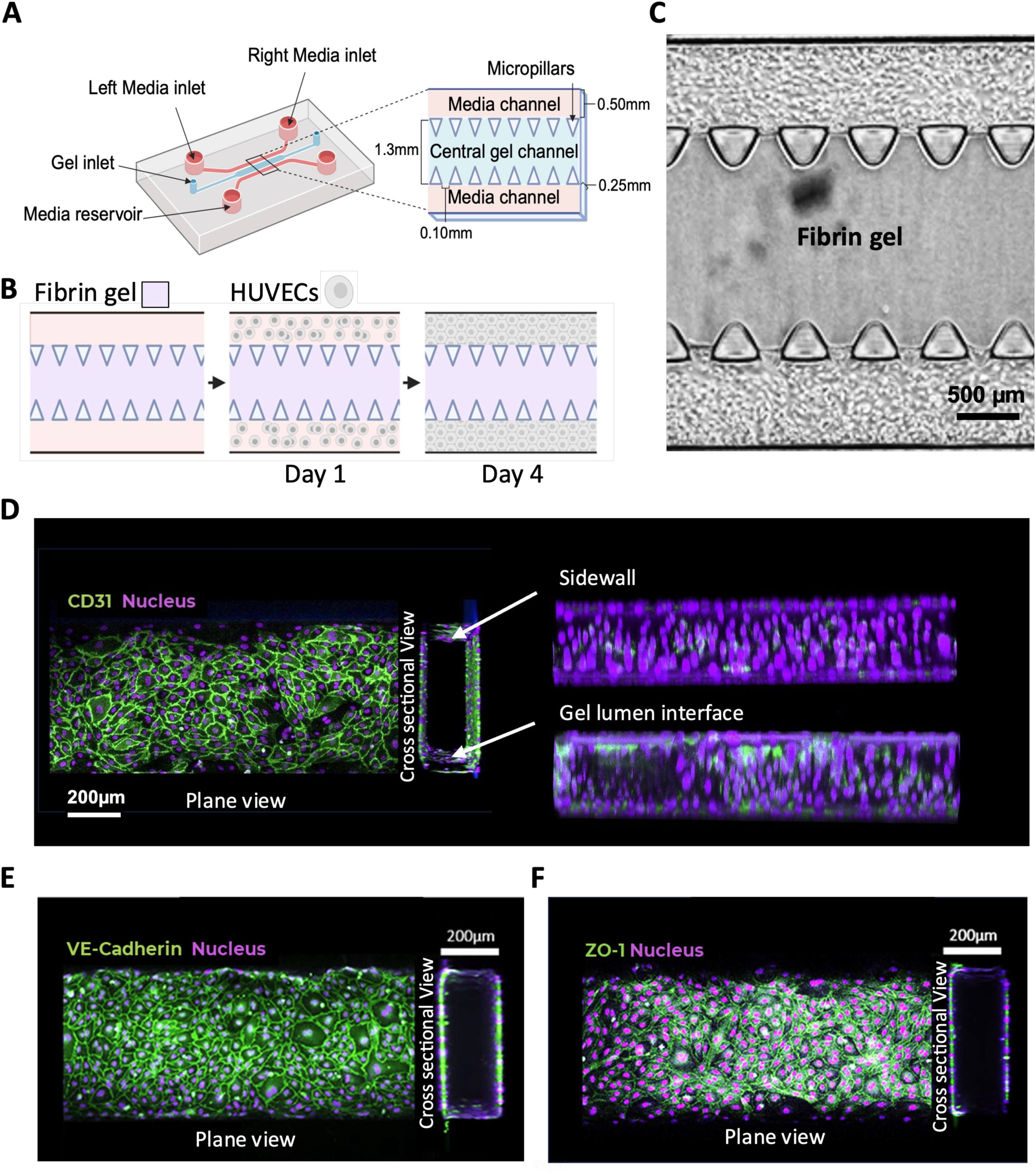
Characterization of parental vessels (PVs) generated in lateral media channels. A) Schematic diagram of the idenTX microfluidic chip from AIM Biotech. The device comprises three parallel microchannels. The central gel channel is flanked by two lateral media channels. All channels are approximately 10.5 mm long and 0.25 mm high. The central gel channel is separated from the lateral media channels by arrays of trapezoidal micropillars. The chip contains six inlets: four inlets connected to reservoirs feeding the two lateral media channels (two per channel) and two inlets connected to the central gel channel without reservoirs. B) Schematic of the procedure used to generate PVs in the microfluidic chip. The central gel channel was filled with fibrin gel. The two lateral channels were coated with collagen and sequentially seeded four times with 10 µL of HUVECs (3 × 10⁶ cells/mL). PVs were generated within 4 days. C) Brightfield microscopy image of PVs formed in the lateral media channels with fibrin gel in the central gel channel. D-F) Representative confocal microscopy images showing planar view of the top surface of the lateral media channel and cross-sectional views of HUVECs stained for CD31 (D), VE-Cadherin (E) or ZO-1 (F). Purple: Nuclei.

### Fibrin hydrogel preparation

For all experiments, fibrin hydrogels were prepared using fibrinogen (6 mg/mL; Sigma-Aldrich, cat. no. 9001-32-5) dissolved in PBS. Fibrinogen solutions were aliquoted, stored at −80 °C and thawed as needed. Thrombin (Sigma-Aldrich, cat. no. 9002-04-4) was initially reconstituted in PBS to a stock concentration of 50 U/mL, aliquoted, and stored at −80 °C. Prior to gel preparation, thrombin was thawed and diluted to a final concentration of 4 U/mL in EBM-2 supplemented with 200 IU/mL penicillin and 0.2 mg/mL streptomycin. Equal volumes of fibrinogen and thrombin solutions were mixed immediately before use, and subsequent steps were performed without delay to prevent premature gelation.

### Generation of Parental vessel in the two lateral media channels

To generate a complete 3D endothelial lining within each lateral media channel, 10 µL of fibrin hydrogel (1:1 mixture of fibrinogen and thrombin) was injected into the central gel channel of idenTX chips (AIM Biotech, cat. no. DAX-1) and allowed to polymerize at 37 °C for 15 minutes. Both lateral media channels were then coated with 0.1 mg/mL type I collagen (Corning, cat. no. 354236) for 15 minutes and rinsed once with supplemented EBM2 medium. HUVECs were suspended at 3 × 10⁶ cells/mL in supplemented EBM2, and 12 µL was injected into the left inlet of the top lateral media channel. The chip was kept vertically (width of the chip 90° relative to the culture surface) for 1 minute, allowing cells to settle against the gel-channel interface through gravity. The chip was then placed face-down in the incubator for 10 minutes to promote cell adhesion. The remaining surfaces (side wall opposite the gel, top wall, and bottom wall) were sequentially coated using additional 12 µL injections with defined chip orientations. For the side wall opposite the gel, a second 12 µL injection was performed, followed by vertical orientation in the opposite direction for 1 minute. For the top wall, another 12 µL injection was performed, and the chip was incubated face-down for 10 minutes to seed the upper surface. Finally for the bottom wall, a 12 µL injection was followed by a 10-minute face-up incubation to seed the lower surface. This multi-orientation seeding strategy ensures that all four internal surfaces of each lateral media channel are uniformly lined with HUVECs. The same procedure was repeated for the opposite lateral media channel. After the final seeding steps, the chip was incubated for 15 minutes at 37 °C. Then, 50 µL of supplemented EBM2 was added to each media reservoir. Chips were maintained at 37 °C with daily medium changes until HUVECs established a continuous, confluent 3D monolayered tube called the Parental Vessel (PV).

All medium changes were performed by aspirating the entire medium volume from the reservoir wells connected to the two lateral media channels of the chip and immediately refilling with fresh supplemented EBM-2 (50 µL in the right reservoir and 70 µL in the left reservoir of each lateral media channel). After medium change, all reservoirs were emptied again and refilled with 50 µL of medium. An additional 20 µL was added to the top-left reservoir of the upper lateral media channel to promote gravity-driven flow across the channels. Care was taken to avoid introducing air bubbles at the inlets of the lateral media channels. Medium was replaced every 24 hours for all chip cultures unless otherwise specified.

To evaluate the integrity of the endothelial barrier, permeability of 70 kDa FITC-dextran was assessed. On day 5 after making the PVs, cell culture medium was aspirated from the media reservoir of both the upper and lower lateral media channels. The chip was placed on the stage of a Nikon A1 HD25/A1R HD25 confocal microscope, and the imaging plane was set using a 10× objective to focus on the upper lateral media channel containing the PV used for tracer perfusion and permeability analysis. A solution of 70 kDa FITC-dextran (1 mg/mL; Sigma-Aldrich, cat. no. 60842-46-8) prepared in supplemented EBM-2 was added to the right reservoir of the upper lateral media channel (70 µL), while supplemented EBM-2 without tracer was added to the right reservoir of the lower lateral media channel (70 µL). Subsequently, 50 µL of FITC-dextran solution was added to the left reservoir of the upper lateral media channel, and 50 µL of supplemented EBM-2 without tracer was added to the left reservoir of the lower lateral media channel. Time-lapse imaging was initiated immediately, and fluorescence images were acquired every minute for 40 minutes.

For experiments involving angiopoietin-1 (Ang-1; R&D Systems, cat. no. 923-AN/CF), PVs were treated with Ang-1 for 24 hours prior to permeability measurements.

### Generation of Microvascular network connected to the Parental Vessels

Microvascular networks (MVNs) were generated based on a previously published protocol from Wan *et al*., which describes two approaches for seeding cells into the central gel channel (one-step and two-step protocol) (26). Both methods were used in this study for different purposes.

To optimize cell concentrations for our final model, the one-step method was used (Figure S2A). In this approach, HUVECs and hLFs were suspended in EBM2 medium containing thrombin (4 U/mL). Five µL of the cell suspension and 5 µL of fibrinogen (6 mg/mL) were mixed, and 10 µL of the resulting hydrogel-cell mixture was immediately injected into the central gel channel. The chip was incubated at 37 °C for approximately 15 minutes to allow polymerization, followed by the addition of 50 µL of supplemented EBM2 into all the media reservoirs of the two lateral media channels.

For the two-step seeding strategy (Figure 2A and S4A), 10 µL of highly concentrated HUVEC suspension (15 × 10⁶ cells/mL) was first introduced into the central gel channel through the left inlet and aspirated to leave a residual coating in between the microposts. A second injection of 10 µL containing HUVECs (10 × 10⁶ cells/mL), hLFs (1 × 10⁶ cells/mL), and Meso13 cells (0.25 × 10⁶ cells/mL), unless otherwise specified, was introduced through the right inlet of the central gel channel to seed the middle region without disrupting the residual layer coating. The chip was incubated at 37 °C for 15 minutes to allow matrix polymerization.

**Figure 2.**
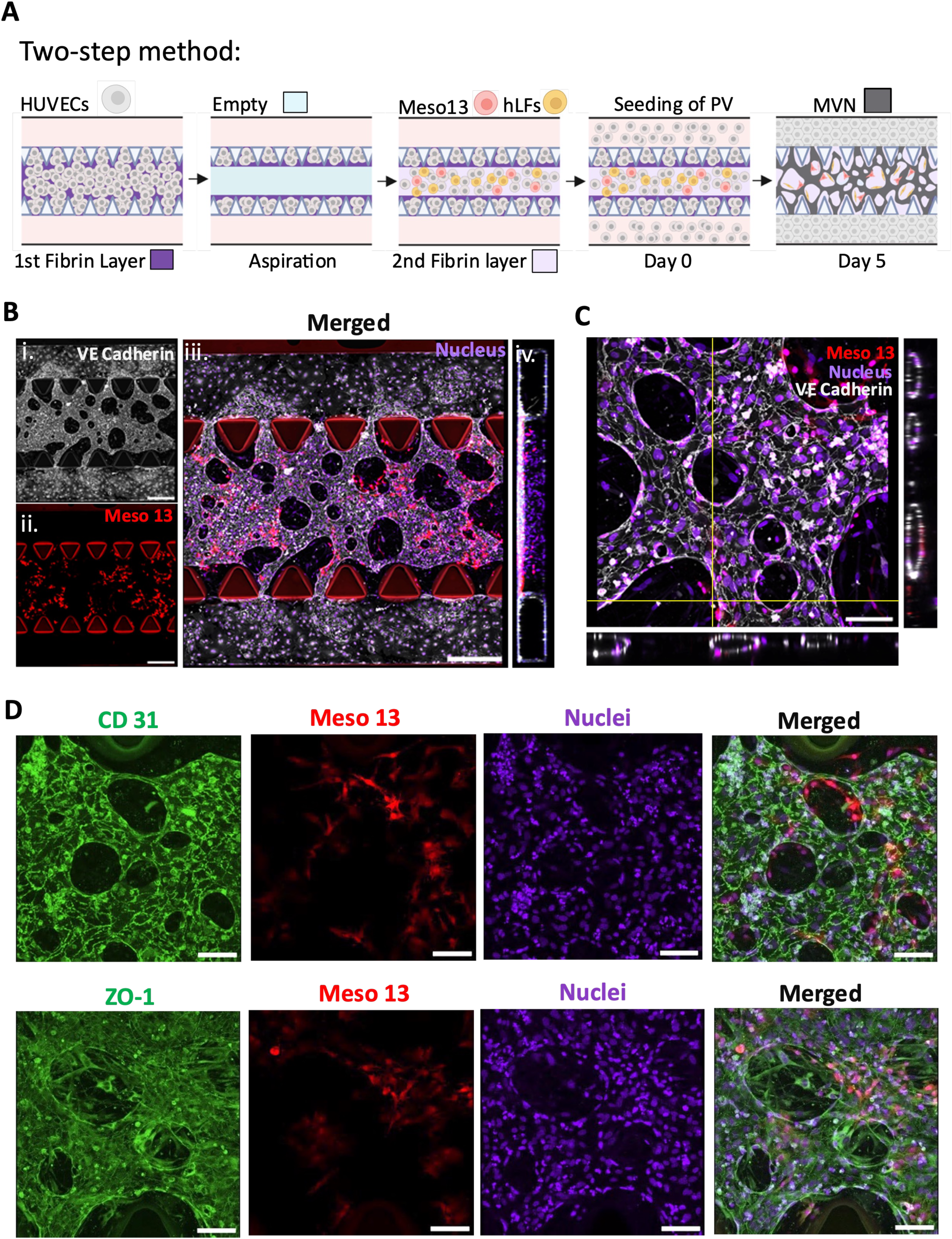
Development of tumor-associated microvascular network (MVN) connected to Parental Vessels (PV) in the chip. A) Schematic of the procedure used to obtain the full model using a two-step method to form an MVN composed of HUVECs, hLFs, and Meso13-BFP cells, connected to two PVs made of HUVECs. The two-step method consists of injecting 10 µL of fibrin gel containing 15 × 10⁴ HUVECs into the central gel channel from the left inlet and immediately aspirating it, leaving a thin layer of cells trapped in the micropillar gaps. A second fibrin gel layer is then seeded from the right inlet by adding 10 µL of fibrin gel containing a suspension of HUVECs (10 × 10⁴ cells), hLFs (1 × 10⁴ cells), and Meso13-BFP cells (0.25 × 10⁴ cells). After complete polymerization of the gel, the PVs are formed by seeding HUVECs in the two lateral media channels. B) 3D projection of confocal images showing VE-cadherin immunostaining (grey) (i), Meso13-BFP (red) (ii), and a merged image of VE-cadherin, Meso13-BFP, and nuclei (purple) (iii), with (iv) an orthogonal view of the chip on day 5 post seeding. Scale bar = 500 µm. C) Orthogonal view showing the presence of hollow lumen in the MVN in the central gel channel of (B). Scale bar =100 µm. D) Representative confocal microscopy images of the MVN after CD31 and ZO-1 labeling (green). Red: Meso13-BFP; purple: nuclei. Scale bar = 100 µm.

To establish lateral connections between the MVN that will form in the central gel channel and the lateral media channels, PVs were generated in both the lateral media channels using the multi-orientation seeding protocol described above, which can be applied directly here as the central gel channel was already filled with polymerized gel. Chips were maintained at 37 °C with 5 % CO₂ and daily medium changes. MVNs were formed, stabilized and connected to the PVs by day 5 post-seeding, at which point the vascularized mesothelioma-on-chip (VMOC) model was set for subsequent experiments.

To assess MVN perfusability in the central gel channel, medium was aspirated from the media reservoir of the two lateral media channels and replaced with fresh supplemented EBM-2 (50 µL per well). The chip was placed on the stage of a Nikon A1 HD25/A1R HD25 confocal microscope, and the imaging plane was set using a 10× objective focused on the central gel channel containing the MVNs. Twenty µL of 70-kDa FITC-dextran solution was added to the left media reservoir of the upper lateral media channel. After a delay of 5–10 seconds, 20 µL of FITC-dextran was added to the left reservoir of the lower lateral media channel. The Z-position was adjusted based on the FITC fluorescence signal to ensure optimal focal depth. Time-lapse imaging was initiated immediately after FITC-dextran loading, and images were acquired in the FITC channel every 15 seconds for 40 minutes.

### Immunofluorescence staining and confocal imaging

Fixation was carried out by filling lateral media channels with 4% paraformaldehyde (EM grade, cat. no. GF720170-1010) and incubating for 1 hour. Permeabilization was done with 0.1% Triton X-100 for 5 minutes. Chips were blocked with 4% bovine serum albumin (BSA) at 4°C overnight and washed with PBS. Chips were stained with Alexa Fluor 488 mouse anti-human CD31 (1:500, BioLegend, cat. no. 303110), Alexa Fluor 488 mouse anti-human ZO-1 (1:100, Invitrogen, cat. no. 339188), or rabbit anti-human VE-cadherin (1:1000, Invitrogen, cat. no. PA5-19612). For the unconjugated VE-cadherin antibody, a secondary Alexa Fluor 488 goat anti-rabbit IgG antibody (1:2000, Life Technologies, cat. no. A11034) was added on the following day. DR (1:5000, Thermo Fisher Scientific, cat. no. 62254) was used as a nuclear stain in all conditions. All conjugated, primary, and secondary antibodies were diluted in 2% BSA. After a final set of PBS washes, chips were ready for imaging.

Fluorescence images were captured using Nikon A1 HD25/A1R HD25 confocal microscope with 4×, 10×, or 20× objectives and the NIS-Elements software (Nikon, RRID:SCR_014329). Z-stacks were acquired using step sizes of 30 µm for the 4× objective, 3 µm for the 10× objective, and 1.5 µm for the 20× objective. Images were processed and reconstructed using NIS-Elements (Nikon), Fiji (ImageJ) and Imaris (Bitplane) software. The NIS Denoise.ai module was used for noise reduction to improve visual clarity. All image quantifications were done on raw images.

### Cell collection from the chip

Cells were collected separately from the central gel channel and the two lateral media channels. Lateral media channels were first washed with PBS, followed by the injection of 50µL of TrypLE Select (Gibco, cat. no. 12563-029) into the left inlet of each lateral media channel and incubation at 37°C for 15 minutes. After incubation, gentle up-and-down pipetting through the left inlet of both channels was performed to facilitate cell detachment. Detached cells were collected and neutralized with RPMI medium supplemented with 10% FBS. This collection step was repeated once more to maximize recovery. For recovery of cells in the central gel channel, 50µL of Liberase™ (50 µg/mL; Roche, cat. no. 05401119001) was injected into the left inlet of both lateral media channels and incubated for 15 minutes at 37 °C to digest the fibrin gel. Released cells were collected through the media inlets. All collected cells were kept at 37°C prior to subsequent use.

### Single cell RNA sequencing and analysis

After 5 days of culture, cells from the central gel channel and lateral media channel (with or without mesothelioma cells) were collected as described above. Single-cell suspensions were prepared in supplemented EBM2 by mixing cells from the central gel channel and lateral media channels at a 9:1 ratio. 10,000 cells per sample were used to generate single-cell droplet libraries with Chromium NEXT GEM Single Cell 3’ Reagent kit v3.1, 10X Chromium Single Cell Controller (10X Genomics, Pleasanton, CA, USA). Library size was determined by Agilent TapeStation assays and the concentration by Qubit Flex (Invitrogen). Libraries were pooled and sequenced on NovaSeq 6000 Sequencing System (Illumina), providing a read depth of >40,000 read pairs per cell according to the manufacturer’s instructions. Four samples were sequenced in two batches, each batch being composed of one chip with mesothelioma cells and one chip without.

Gene expression matrices were generated using Cell Ranger (10X Genomics, version 2.1.1) with the GRCh38 build of the human reference genome, and further processed using RStudio (version 2023.12.1+402). We used the following quality control steps: genes expressed by <3 cells were not considered, cells expressing <400 genes (low quality) or >10,000 genes, <500 genes or >80,000 unique molecular identifiers (UMIs), or >15% of UMIs derived from the mitochondrial genome were removed. From the mesenchymal and endothelial clusters, doublets were detected and labelled using the DoubletFinder package (version 2.0.6). Doublets represented ∼0.5% of filtered cells and were evenly distributed among the different cell types from all samples. Notably, doublet annotation incorrectly identified the entire tumor cluster as a doublet, probably due to the active nature of this cell type compared to the other cells from the VMOC. Therefore, this annotation was not considered for quality filtering of the tumor cells. Since enzymatic and mechanical dissociations were described as eventually inducing a specific transcriptomic response, we checked if a specific cell cluster/subcluster was enriched for the dissociation signature (27). Visualization, clustering, marker gene identification, and gene set enrichment analysis (GSEA) were performed using the SeuratV5 package. Signature enrichment scores were calculated using the UCell R package (version 2.12.0) using the Mann-Whitney U statistic. Cell trajectory analysis was performed using the Monocle3 R package (version 1.4.26). CellChat was performed using default parameters on the full CellChat database (version 2.2.0) by predicting interactions between all subclusters composing the VMOC. Gene Set Variation Analysis (GSVA) was performed with the GSVA R package (version 2.2.1) on the top 50 marker genes from each endothelial and mesenchymal subcluster. Deconvolution was achieved on endothelial and mesenchymal subclusters with the MuSiC R package (version 1.0.0) using common genes expressed between scRNA-seq and bulk 3’RNA-seq datasets generated in this study.

### MV-GFP infection of vascularized mesothelioma-on-chip

The live-attenuated Schwarz strain of measles viruses expressing the green fluorescent protein (MV-GFP) was produced as previously described (28). Viral titer was determined using the TCID_50_ method. TCID_50_ quantifies the infectious dose of a virus by determining the dilution at which 50% of cell cultures show infection.

For virus infection, the VMOCs were washed with supplemented EBM2 and then each media reservoir was filled with 50 µL of the same medium. MV-GFP was introduced by preparing a viral suspension containing 10⁶ TCID₅₀ in a total volume of 40 µL supplemented EBM-2, which was added to the left reservoir of the upper lateral media channel. After 6 hours of incubation at 37°C, the virus solution was aspirated, and the lateral media channels were washed twice with fresh supplemented EBM2 medium. Medium was replaced every 24h. Images were acquired 48 h post-infection using a Nikon A1 HD25/A1R HD25 confocal microscope.

### Luciferase release assay to quantity MV-induced PM cell death in VMOC

2,500 Meso13-luc cells were embedded in the 2^nd^ layer of the central gel channel using the two-step method. Chips were infected with MV-GFP on day 5, as described above. Supernatants were collected at 24-, 48-, and 72-hours post-infection, followed by a medium renewal, and stored at -80°C. For each time point, 50 µL of medium was transferred to a 96-well plate and mixed with 50 µL of Nano-Glo® Luciferase Assay substrate (Promega, cat. no. N1120). Luminescence was measured using a Fluoroskan plate reader (Thermo Fisher Scientific). Each time point was normalized to the uninfected condition.

### Chemokine detection by Legendplex multiplex assay

Supernatants from infected and non-infected VMOCs were collected 48 hours after MV-GFP infection for cytokine measurements. LEGENDplex™ Human Inflammation Panel 1 and LEGENDplex™ Human Immune Checkpoint Panel 1 were used to measure IL-4, IL-2, IP-10, IL-1β, TNF-α, MCP-1, IL-17A, IL-6, IL-10, IFN-γ, IL-12p70, IL-8, CD25 (IL-2Rα), 4-1BB, sCD27, B7.2 (CD86), TGF-β1, CTLA-4, PD-L1, PD-L2, PD-1, Tim-3, LAG-3, and Galectin-9 secretions.

### RNA extraction from VMOC

RNA was extracted using the Nucleospin® RNA plus kit (Macherey-Nagel, cat. no. 740990.50). The mRNA quality was analyzed using the Agilent 20100 Bioanalyzer (Agilent) in RNA Nanochips (Agilent).

### 3’ RNAseq analysis

The 3′ RNA sequencing profiling was performed by the GenoBird platform (IRSUN, Nantes, France) using a NovaSeq 6000 (Illumina) and NovaSeq 6000 SP Reagent Kit 100 cycles (Illumina) according to the manufacturer’s protocol. The full procedure is described in a study by Charpentier *et al*. (29). The raw sequence reads were filtered based on quality using FastQC. Adapter sequences were trimmed from the raw sequence reads using Cutadapt. Reads were then aligned to the reference genome (Hg38) using BWA. Differential gene expression were determined using Biomex software version 1.0-5 (30).

### Statistical data

All data were expressed as mean ± SEM, and statistical analyses were performed with GraphPad Prism V.8 (GraphPad Software, Boston, Massachusetts, USA). Statistical tests and significant differences for each figure were mentioned in the legends.

## Results

### Generation and characterization of parental vessels

As in many solid malignancies, the pre-existing primary vasculature is hijacked to ensure blood supply to the tumor mass and enable metastatic dissemination. Neighboring and newly formed tumor-associated secondary vessels exhibit irregular morphology, increased permeability, and heterogeneous perfusion (31,32). To recapitulate these two distinct vessel types in our vascularized mesothelioma-on-chip (VMOC) model, we used the idenTX AIM-Biotech microfluidic chip (Figure 1A), which consists of a central gel channel (CC) flanked by two lateral media channels (LCs). Trapezoidal micropillars separate the channels, while the gaps between micropillars allow molecular exchange and endothelial connections across channels. To reduce experimental complexity and ensure controlled optimization, the two vessel types were optimized and characterized separately, after which both were generated within the same chip to form the complete VMOC.

To generate the primary vascular compartment, i.e., the parental vessel (PV), the CC was first filled with fibrin gel to prevent endothelial cells (ECs) seeded in the LC from spreading into the CC. Because PV formation requires multiple injections through the LCs (Figure 1B), the extracellular matrix (ECM) in the CC needs to remain mechanically stable throughout the process. Fibrin was selected over collagen because it is less deformable under repeated injections (33) and maintains strong adhesion to the micropillars. The polymerization process was optimized to ensure a stable and reproducible gel. A fibrinogen-thrombin mixture (6 mg/mL and 4 NIHU/mL, respectively) was selected for all experiments, injected into the CC and allowed to polymerize at 37°C for 15 minutes. This incubation period was the most effective, as an extended duration resulted in gel contraction or the formation of air bubbles between the micropillars. After fibrin polymerization, LCs were coated with collagen to promote cell adhesion, and HUVECs were injected with sequential 90° rotations to ensure uniform attachment of cells along all four walls of both media channels (Figure 1B). After 4 days, HUVECs formed a 3D continuous cell monolayer covering the entire LC (Figure 1C and 1D).

Confocal microscopy showed the monolayer of CD31+ HUVECs all around the inner surface of the LCs (plane view) and the presence of a hollow lumen (cross-sectional view) (Figure 1D). This resulting tube-like structure is referred to as the PV. Immunofluorescence labeling for VE-cadherin and ZO-1 demonstrated the presence of adherens and tight junctions, respectively (Figure 1E and 1F). In order to evaluate the response of the blood vessels to external stimuli, the barrier functionality of the PV was evaluated using a 70 kDa FITC-dextran vascular permeability assay (Figure S1A). Quantitative analysis of FITC-dextran diffusion was used to calculate a permeability coefficient (Pd) equal to 8.10 x 10^-6^ ± 4.00 x 10^-6^ cm/s in the presence of HUVEC whereas Pd increased to 1.40 x 10^-5^ ± 1.30 x 10^-6^ cm/s without PVs (Figure S1A). To evaluate the responsiveness of PVs to external stimuli, we used angiopoietin-1 (Ang-1). Ang-1 is a cytokine that promotes maturation and stabilization of blood vessels by stabilizing endothelial junctions through various mechanisms, thereby reducing vascular permeability (34–36). In a previous vasculature-on-chip model, Ang-1 led to a dose-dependent decrease in permeability to FITC-dextran and nanoparticles (37). Consistently, in our model, treatment of PVs with 100 ng/mL Ang-1 for 24 hours resulted in a reduction in 70 kDa FITC-dextran diffusion into the surrounding fibrin gel (Pd = 3.60 x 10^-6^ ± 4.00 x 10^-7^) compared to the untreated condition (Pd = 8.10 x 10^-6^ ± 4.00 x 10^-6^ cm/s) (Figure S1A). The second external stimulus evaluated was the application of a flow of cell culture medium. Fluid flow at 5 µL/minute (a velocity of ∼670 µm/second) for 24 hours induced an alignment of HUVECs in the direction of flow (Figure S1B). This velocity corresponds to the physiological range of capillary blood flow (≈500–1,500 µm/second) (36–38). Overall, these results demonstrated that PVs form a functional endothelial barrier capable of responding to both molecular and mechanical signals.

### Formation of a vascularized tumor microenvironment connected to PV

Tumor-associated vasculature is highly heterogeneous, with irregular branching and architecture, so we aimed to recreate this complex vascular network as our secondary endothelial compartment (38). A one-step seeding approach was first evaluated to establish the secondary endothelial compartment by introducing a HUVEC–fibrin mixture into the CC on day 0 (Figure S2A). Previous studies have shown that HUVECs suspended in fibrin gel undergo vasculogenesis and self-organize into a microvascular network (MVN) (39–41). Primary HUVECs expressing mRuby2 were injected into the CC, allowing visualization of endothelial connections and network formation over time. Because the LCs serve as the primary source of culture medium for the chip, perfusion of the MVN requires direct endothelial connectivity across the micropillar gaps between the MVN in the CC and the PV in the LCs. These endothelial connections, designated as bridging vessels, span the micropillar gaps and form the sole fluidic connection between the PVs and the MVN (Figure S3A-B). An increase in HUVEC seeding density led to larger vessel diameters in the MVN (Figure S2B-D and S3C) (26). At low density (80,000 HUVECs per CC), the number of perfusable bridging vessels was low due to insufficient HUVEC deposition between the micropillar gaps (Figure S2E). Increasing density (80,000–150,000 HUVECs per CC) increased the number of perfusable bridging vessels and improved connectivity with the LCs (Figure S2E). However, high densities (120,000–150,000 HUVECs per CC) produced large, simplified vessels in the central gel (Figure S2B). Overall, this one-step seeding approach did not allow simultaneous formation of a MVN with interconnected vessel structures and sufficient perfusable bridging vessels.

To overcome this limitation, we implemented a two-step seeding strategy adapted from Wan *et al*. (26) (Figure S4A). This approach allows independent control of the region in the micropillar gaps and middle region of the CC. Briefly, two sequential injections of HUVECs in fibrin-thrombin gel were performed. The first with a high concentration of HUVEC to fill the micropillar gaps and the second with a lower HUVEC concentration to fill the middle region of the CC. Once polymerization was complete, PVs were established in both LCs, enabling the PVs to act as macrovessels connected to bridging vessels and supplying medium to the fibrin embedded MVN in the CC. By day 5, the MVN was stable and perfusable (Figure S4B-panel i).

Having established a perfusable and branched secondary MVN that was connected to the PVs, we next introduced stromal cells. Fibroblasts are major components of the TME (42–44) and also play an important function in the structure and formation of vessels (45–47). Thus, to better reflect the stromal context of PM tumors, we incorporated primary human lung fibroblasts (hLFs) into the CC along with the MVN. To do this, we first evaluated the impact of hLFs on the formation of MVN by co-seeding HUVECs with hLFs in the second layer of the two-step method at different HUVECs/hLF ratio (H/F ratio) (Figure S4A-B). Fibroblasts were excluded from the first gel layer to prevent narrowing of the bridging vessels, as previously reported (39). Figure S4B shows that in the absence of fibroblasts (Figure S4B, panel i) and with a H/F ratio = 10:2 (Figure S4B, panel iii), vessels were either less branched or poorly structured. With an H/F ratio = 10:1 (Figure S4B, panel ii), branched, well-formed, and interconnected microvessels were obtained. The perfusion of 70 kDa FITC-dextran injected into the LCs demonstrated the presence of numerous connections between PVs and the MVN in the absence of hLFs and with the H/F ratio = 10:1 (Figure S4B, panel i and ii, respectively), whereas the number of connections was reduced with an H/F ratio = 10:2 (Figure S4B, panel iii). A decrease in lumen size was observed with the increase in the number of hLFs (Figure S4C). This observation was confirmed by a high prevalence of small vessels (50 µm in diameter) and the absence of large vessels (>300 µm) with H/F ratio = 10:1 and 10:2 (Figure S4D). Considering the results obtained, the H/F ratio = 10:1 was used for all the subsequent experiments. In Figure S5A, images show the evolution of the model over a period of 10 days. On day 0, HUVECs were introduced through the left inlet of the CC and were evenly distributed as single cell suspension in the fibrin gel. By day 2, connections between cells appeared. On day 4, network-like structures were visible, though still exhibiting parts with active sprouting and anastomosis. By day 5 and 6, the networks appeared well connected with smoother structures and remained stable up to day 10. To complete the characterization of the MVN, we performed CD31, VE-cadherin and ZO-1 labeling and confirmed the endothelium integrity and presence of adherens and tight junctions (Figure S5B).

Finally, to introduce the tumor compartment of the VMOC, we used the two-step method and added the Meso13 PM cell line to the mix of HUVECs and hLFs (H/F/Meso13 ratio = 10:1:0.25) in the second fibrin gel layer introduced into the CC (Figure 2A). The addition of PM cells did not modify the structure of the MVN, and by day 5, a stable MVN was formed, bridging vessels were preserved, and sufficient cancer cells were present for subsequent experiments (Figure 2B). Immunostaining and confocal microscopy of VE-cadherin showed the presence of adherens junctions in the MVN connected to the PVs (Figure 2B, panel i, iii and iv). Orthogonal projection of confocal microscopy data obtained after Z-acquisition confirmed the presence of round vessels with a hollow lumen in the MVN (Figure 2C). As in Figure S5B, in the presence of PM cells, we observed the presence of CD31 and ZO-1 positive cells (Figure 2D).

To examine the 3D organization of the MVN and its spatial relationship with Meso13 pleural mesothelioma cells, we generated a 3D representation of the MVN with VE-cadherin staining (Figure 3A, panel i), a magnification of an area of the MVN (Figure 3A, panel ii) and a transverse section of the MVN showing a hollow circular lumen (Figure 3A, panel iii). Meso13 cells were distributed throughout the fibrin matrix and were frequently located adjacent to the vessel structures organized into highly branched, complex networks (Figure 3A, panel i and iii and Movie S1). To demonstrate functionality and interconnection between the MVN and PVs, a perfusion test using 1 µm fluorescent microbeads was performed on day 5 of culture. After addition of microbeads to the media reservoir of the upper LC, beads traveled from one PV through the MVN to the opposite PV, demonstrating perfusability and continuous luminal connectivity between vascular compartments of the LCs and CC (Figure 3C and Movie S2). In addition, by overlaying the bead signals with the fluorescent signals from the vessels, 3D reconstructions confirmed that the microbeads were localized within the MVN lumens (Figure 3B). Perfusion of 70-kDa FITC–dextran through the MVN also clearly visualized interconnected and perfusable vessels spanning the central gel region (Figure 3D).

**Figure 3.**
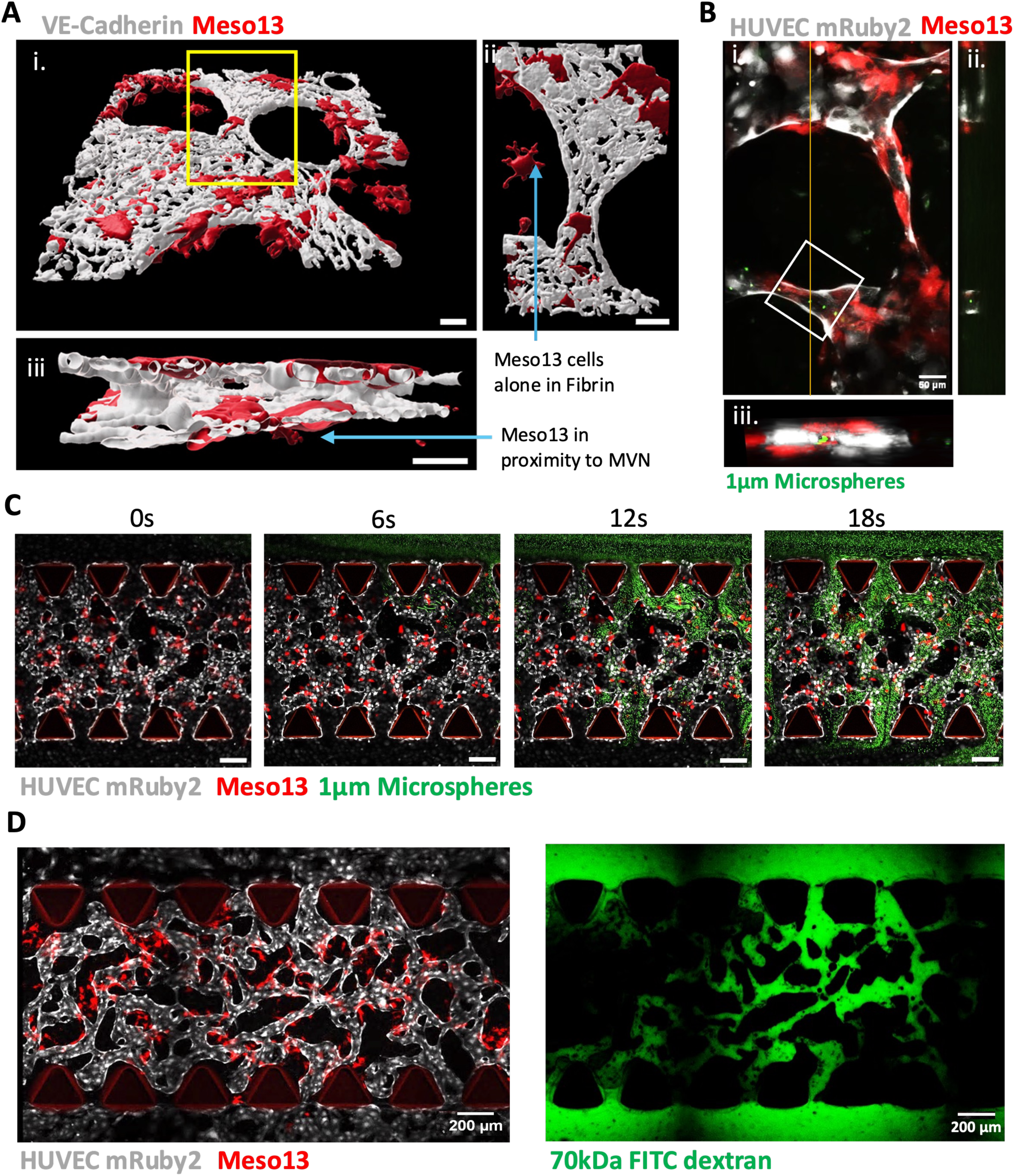
Characterization of the microvascular network (MVN) by 3D reconstruction, and perfusion with 1 µm fluorescent microbeads and 70 kDa FITC-dextran. A) Confocal imaging and 3D Imaris reconstruction of the MVN. (i) Maximum intensity projection showing the MVN. (ii) Planar view of the area defined by the yellow rectangle in (i). iii) Transversal view of the area defined by the yellow rectangle in (i). White: VE-cadherin; red: Meso13-BFP. Scale bar = 50 µm. B) Confocal microscopy images and orthogonal projections showing 1 µm green-fluorescent polystyrene beads (microbeads) within a vessel of the MVN. ii) Orthogonal projection of a 3D stack showing microbeads in the lumen of the MVN, and (iii) 3D image reconstruction of the white square in (i) showing microbeads in the MVN. C) Diffusion of microbeads in the VMOC over time after injection into one parental vessel (PV). Scale bar = 200 µm. D) Confocal microscopy images. Left panel: structure of the MVN in the central gel channel. Grey: HUVECs-mRuby2; red: Meso13-BFP. Right panel: diffusion of 70 kDa FITC-dextran into the same MVN (green).

### Evaluation of cellular heterogeneity in the VMOC model

A recurrent question in the development of tumor models *in vitro* is the extent of cellular heterogeneity. Indeed, it is now well established that cells from the TME exist under different subpopulations harboring distinct properties (48,49). Seemingly, the VMOC clearly shows a heterogeneity, with different vessel organization in PVs and MVN. To evaluate this heterogeneity, we performed single-cell RNA-sequencing (scRNA-seq) analysis on VMOCs with or without Meso13 cells after 5 days of culture. After quality filtering, doublet removal and integration, we obtained 37,215 high-quality cells that grouped into three distinct clusters (Figure 4A). Annotated on the basis of their top 50 marker genes, we retrieved the three cell types included in the VMOC, namely cancer cells, fibroblast-like (mesenchymal) and endothelial cells, with a homogeneous distribution across the 4 samples (Figure 4A-B, S6A and Table S1). Considering the initial cell ratio used in the VMOC, the cancer cell subset was constituted of 318 cells that fell into three subclusters with a similar relative proportion across samples (Figure 4C and S6B-C). As expected, no cancer cells were detected in the samples devoid of Meso13 cells (Figure S6A). Gene set enrichment analysis (GSEA) showed enrichment in specific transcriptomic signatures including stress and inflammatory responses, oxidative phosphorylation (OXPHOS) and cell proliferation (Figure 4C, S6D-F, Table S1). Interestingly, CellChat analysis identified a TGF-β signaling network with multiple inferred ligand–receptor interactions, with cancer cells as the main signal senders to endothelial cells and fibroblast (Figure S9A, panel i and ii).

**Figure 4.**
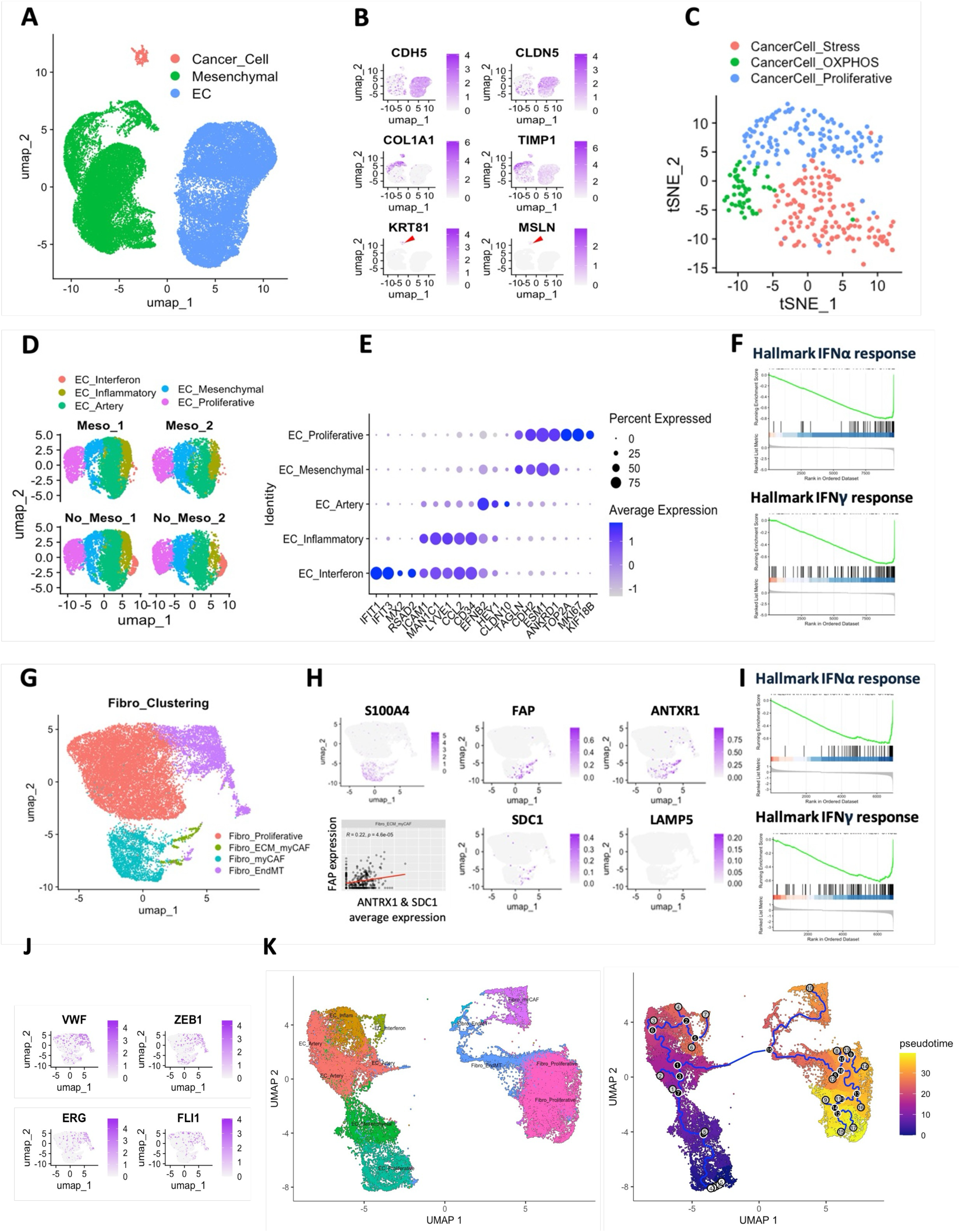
Characterization of the VMOC by single-cell RNA sequencing (scRNA-seq). A) UMAP representation of cell types from the VMOC following scRNA-seq (n = 2), including pleural mesothelioma (PM) cells, fibroblast-like (mesenchymal) cells and endothelial cells, B) Feature plots of all cell types in VMOC with different marker gene signature, identifying endothelial cells (*CDH5*, *CLDN5*), fibroblast-like/mesenchymal (*COL1A1*, *TIMP1*), and PM cells (*KRT8*, *MSLN*). C) t-SNE projection of PM cells showing segregation into transcriptionally distinct states. D) UMAP projections of endothelial cells from VMOC conditions with PM cells (Meso; n = 2) and without PM cells (No_Meso; n = 2). E) Dot plot showing the expression of selected marker genes across endothelial cell subpopulations. Dot size represents the percentage of cells expressing each gene, and color intensity indicates the average expression level. F) Enrichment plots of hallmark interferon-α and interferon-γ response gene sets in endothelial cells from VMOC conditions with PM cells compared to conditions without PM cells. G) UMAP representation of fibroblasts from the VMOC showing clustering into distinct transcriptional states H) and marker genes for CAF subtypes (*S100A4*, *FAP*, *ANTXR1*, *SDC1*, and *LAMP5*). Scatter plot (bottom left) shows the relationship between FAP expression and the average expression of *ANTXR1* and *SDC1* for Fibro_ECM_myCAF subtype. I) Enrichment plots of hallmark interferon-α and interferon-γ response gene sets in fibroblasts from VMOC conditions with PM cells compared to conditions without PM cells. J) Feature plots of fibroblasts with marker genes for endothelial identity and endothelial-to-mesenchymal transition (EndMT). K) UMAP and pseudotime representation of endothelial and fibroblast populations. Cells are colored by annotated transcriptional states (left panel) and by pseudotime values (right panel), with the inferred trajectory overlaid.

Thereafter, we sought to investigate how cancer cells impact on the vascular and mesenchymal compartments at the single cell level. The 18,151 endothelial cells obtained post-filtering segregated into 5 distinct clusters with a similar distribution across samples (Figure 4D and S7A-B). We could first identify two populations previously described in scRNA-seq datasets from cultured endothelial cells (50,51), namely proliferative endothelial cells expressing key proliferation markers (*MKI67*, *TOP2A*, *BIRC5*) and genes related to chromosomal segregation, and a mesenchymal transitory population (Figure 4D-E, S7C-E and Table S1). Interestingly, a subcluster of endothelial cells expressed markers associated with artery specification (*HEY1*, *EFNB2*, *CLDN10*) (52) that was absent in scRNA-seq datasets from static 2D culture (Figure 4E) (50,51). Hence, suggesting that a vascular specification program may occur in the VMOC, in response to 3D architecture. Consistently, CellChat analysis revealed a strong NOTCH signaling network between endothelial subpopulations, with multiple ligand–receptor interactions, including *DLL4*–*NOTCH1*, supporting its role in arterial fate specification (Figure S9B, panel i and ii) (53–55). The last two endothelial subclusters expressed high levels of genes related to pro-inflammatory cytokines and response thereof (*CCL2*, *CXCR4*, *ICAM1*, *LYVE1*, *EGR1-3, NUPR1*), but also key angiogenic markers (*KDR*, *PGF*, *DLL4*) (Figure 4D-E and Table S1). Interestingly, among the inflammatory subclusters, one subset coined “EC_Interferon” presented a marked signature in interferon signaling (with key interferon-stimulated genes (ISGs): *IFIT1-3*, *IFIT5*, *IFI6*, *IFI44L*, *MX1-2*, etc.) (Figure 4D-E, S7F-G and Table S1), and its abundance was reduced by 95% when mesothelioma cells were added into the VMOC (Figure 4D and S7B). In addition, a decrease in genes associated with interferon alpha and gamma response in the whole EC population was also observed following the addition of PM cells (Figure 4F and S7H).

In the VMOC model, the fibroblast compartment was abundantly represented with 18,746 high quality cells, distributed in four different subclusters (Figure 4G and S8A-B). The largest fibroblast subcluster expressed genes and gene sets associated with cell proliferation, chromatin remodeling and nucleosome formation (*PCLAF*, HMGB2, *H2AFZ*) but lacked expression of the cancer-associated fibroblast (CAF) marker FSP1 (*S100A4*) (Figure 4H). On the contrary, two fibroblast subclusters expressed CAF markers, characteristic of myofibroblasts (myCAF) (56) and ECM-myCAF (*FAP*^+^ *ANTXR1*^+^ *SDC1*^+^ *LAMP*^−^) (Figure 4H and Table S1). Interestingly, a final fibroblast subcluster coexpressed endothelial (*CDH5*, *PECAM1*, *VWF*, *CLDN5*) and mesenchymal markers, with key transcription factors involved during the endothelial-to-mesenchymal transition (EndMT) including *ZEB1*, *ERG* and *FLI1* (Figure 4J and Table S1) (57). Monocle cell trajectory analysis revealed that this fibroblast subset (named Fibro_EndMT) may originate from an ultimate differentiation of endothelial cells into fibroblasts via EndMT (Figure 4K and S8C). Finally, mirroring the effect seen in endothelial cells, the addition of mesothelioma cells in the VMOC induced a sharp diminution in gene sets associated with IFN alpha and gamma response, and TNFα signaling in the fibroblast subsets (Figure 4I, and S8D).

### Establishment of MV infection and assessment of oncolytic activity in the VMOC

We have previously shown that human PM cell lines intrinsically deficient in IFN-I production and sensitive to MV infection, such as Meso13, still retain functional IFN-α/β receptor (IFNAR) signaling and can be protected by other cells from the TME through paracrine production of IFN-I (17). To better understand the mechanism of sensitivity/resistance to MV, models integrating tumor cells and cells from the TME are necessary. We studied replication, spread, and oncolytic activity of the live-attenuated, vaccinal Schwarz strain of MV coding for GFP (MV-GFP) in the VMOC. We evaluated multiple delivery strategies and viral doses to define the best conditions for studying the MV mechanism of action. MV added directly to the CC during gel loading at multiplicities of infection (MOI; ratio of infectious viral particles to target cells) of 1 and 10, or pre-infection of Meso13 cells at an MOI of 1 prior to embedding, resulted in low PM cell infection, with GFP detected mainly in fibroblasts (Supplementary Figure S10).

We then assessed intravascular administration by introducing the virus through the media inlet of the LC containing the PV after VMOC maturation (day 5) (Figure 5A).

**Figure 5.**
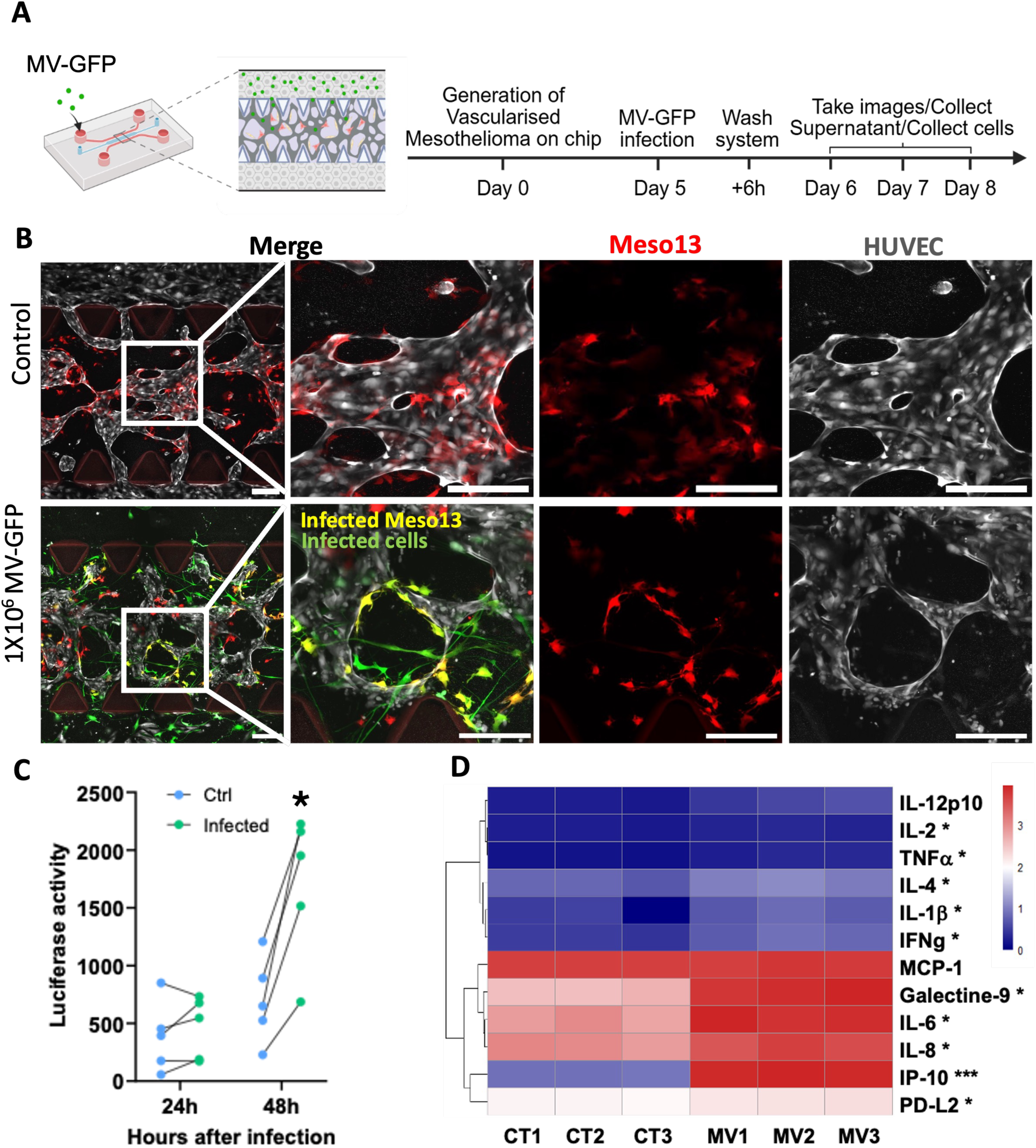
MV infection in the microfluidic chip induces tumor cell death and triggers secretion of pro-inflammatory cytokines. A) Schematic of the MV infection protocol in the chip. 1x10^6^ MV-GFP particles were injected into a PV on day 5 of the MVN formation. After 6h, MV-GFP was removed, and analyses were performed at 24h, 48h and/or 72h post infection. B) Representative confocal microscopy images of MV-GFP infection at 48h post infection. Green: GFP expressing cells (Infected cells); red: Meso13-BFP; yellow: Infected Meso13; grey: HUVEC. GFP signal detected in regions lacking overlap with both PM (red) and HUVEC (grey) markers was considered indicative of infected fibroblasts. Scale bar = 200 µm. C) Nanoluciferase release from Meso13-luc cells after MV-GFP infection. Supernatants were collected at 24- and 48-h post infection and nanoluciferase activity was measured. n=5, *p<0.05, Unpaired t-test (two-tailed). D) Heatmap of cytokine concentrations measured in samples collected at 48h post infection using LEGENDplex Assay. Representations were obtained using log10 values. n=3, *, p<0.05; **, p<0.01; ***, p<0.001. Unpaired t-test (two-tailed).

Several combinations of viral concentration and exposure duration were tested to identify the optimal conditions for PM cell infection. Preliminary experiments showed that the system could not tolerate stagnant medium for more than 24 hours, as absence of medium renewal resulted in MVN collapse (Figure S11A). Based on this, mature VMOCs were first exposed to 0.2x10^6^, 0.5x10^6^, or 1x10^6^ TCID_50_/viral particles for 24 hours. At 48 hours after infection, 0.2x10^6^ viral particles produced minimal infection but retained vascular integrity (Figure S11B). In contrast, 0.5x10^6^ and 1x10^6^ viral particles produced numerous detectable GFP signals but caused pronounced disruptions and complete collapse of the MVN (Figure S11B). We then tested a high-dose, short-exposure strategy. 1x10^6^ and 2x10^6^ viral particles of MV-GFP were added in a PV for 6 hours and then removed (Figure 5A). Viral replication, as indicated by GFP signal, was detectable as early as 24 hours post-infection with 1x10^6^ viral particles (Figure S11C). Administration of 1x10^6^ viral particles in a PV led to the infection of numerous PM cells and of some fibroblasts at 48 hours after infection (Figure 5B). With 2x10^6^ viral particles, the number of infected PM cells increased as well as the number of infected fibroblasts and an excessive alteration of the MVN was observed, despite low infection of endothelial cells (Figure S11D). Therefore, the condition of 1 × 10⁶ viral particles and 6 hours of incubation was selected as the optimal condition for establishing efficient PM cell infection.

In order to specifically measure the death of PM cells, we used Meso13 cells expressing nanoluciferase (Meso13-luc). Detection of luciferase activity in the culture supernatant is proportional to the extent of PM cell death. We observed a basal nanoluciferase activity in control conditions that remained stable between 24 and 48 hours. In contrast, MV-GFP administration increased nanoluciferase activity relative to control, indicating virus-induced lysis of PM cells (Figure 5C). PM cell death increased as early as 24 hours post-infection and became significant at 48 hours after infection. These data demonstrate that MV induced measurable and significant lysis of PM cells in the VMOC after 48 hours of infection.

### MV infection induces inflammatory response in the VMOC

MV infection is well documented to trigger robust innate immune activation, including induction of ISGs, as well as production of inflammatory chemokines and cytokines (58–60). To study the consequences of MV infection on the VMOC, inflammatory cytokines were measured in culture supernatants 48 hours post-infection using multiplex ELISA (Figure 5D and S12). MV infection induced significant release of numerous cytokines compared to the non-infected controls. IP-10 (CXCL10) showed the strongest induction (>1,000-fold), followed by substantial increases in IL-6, IL-8, Galectin-9, TNF-α and IFN-γ, indicating activation of both interferon-responsive and inflammatory pathways. In contrast, IL-12p70 and MCP-1 remained unchanged, possibly due to their already high basal expression in control condition, particularly for MCP-1.

To further characterize the transcriptomic programs induced following MV infection, we performed a 3’RNA-seq analysis on the whole VMOC chips at 24- and 48-hours post-infection (Figure 6). MV infection induced extensive transcriptional modifications in the VMOC, with 197 genes upregulated and 2 downregulated at 24 hours, and 210 upregulated and 25 downregulated at 48 hours (padj < 0.01) (Figure 6A and Table S2). The majority of the deregulated genes are common between the two time-points (n=157) (Figure S13A and Table S2). Pathway enrichment analysis revealed strong activation of innate immune and inflammatory pathways, including type I and type II interferon responses and inflammatory signaling at both time points (Figure 6B and S13B). Figure 6C shows that ISGs were strongly upregulated at both time points. Genes described as hallmarks of response, resistance and toxicity to immune checkpoint blockade (ICB) were also affected following MV infection, with an induction of genes involved in antigen presentation, *HLA-A*, *HLA-B*, *HLA-C* and *B2M*, but also in immune response inhibition, *CD274* (encoding PD-L1), *CD47,* and *HLA-E* (Figure 6D). Endothelial activation markers, such as *SELL*, *SELE*, *VCAM1*, and *ICAM1*, were also upregulated after MV infection, indicating that the vascular compartment was impacted by MV and contributed to the inflammatory response (Figure 6E). In parallel, MV triggered a strong inflammatory transcriptional program by inducing the expression of numerous genes coding for chemokines and cytokines including those previously measured by ELISA in the culture medium, *CXCL8* (IL-8), *LGALS9* (Galectin-9), *CXCL10* (IP-10) and *IL6* (Figure 6F). Finally, we applied deconvolution and gene set variation analysis (GSVA) approaches to infer how MV impacts the endothelial and fibroblast subpopulations identified by scRNA-seq (Figure S14A-D). Inflammatory and proliferative endothelial cells increased at 48 hours post MV infection, while this tendency was inverted in proliferative fibroblasts (Figure S14A-B). GSVA from the top 50 marker genes showed increased pathway activity scores from the EC_interferon in all conditions with MV infection (Figure S14C). Altogether these results show that MV induces death of PM cells, impacts the TME and induces strong inflammation in the VMOC.

**Figure 6.**
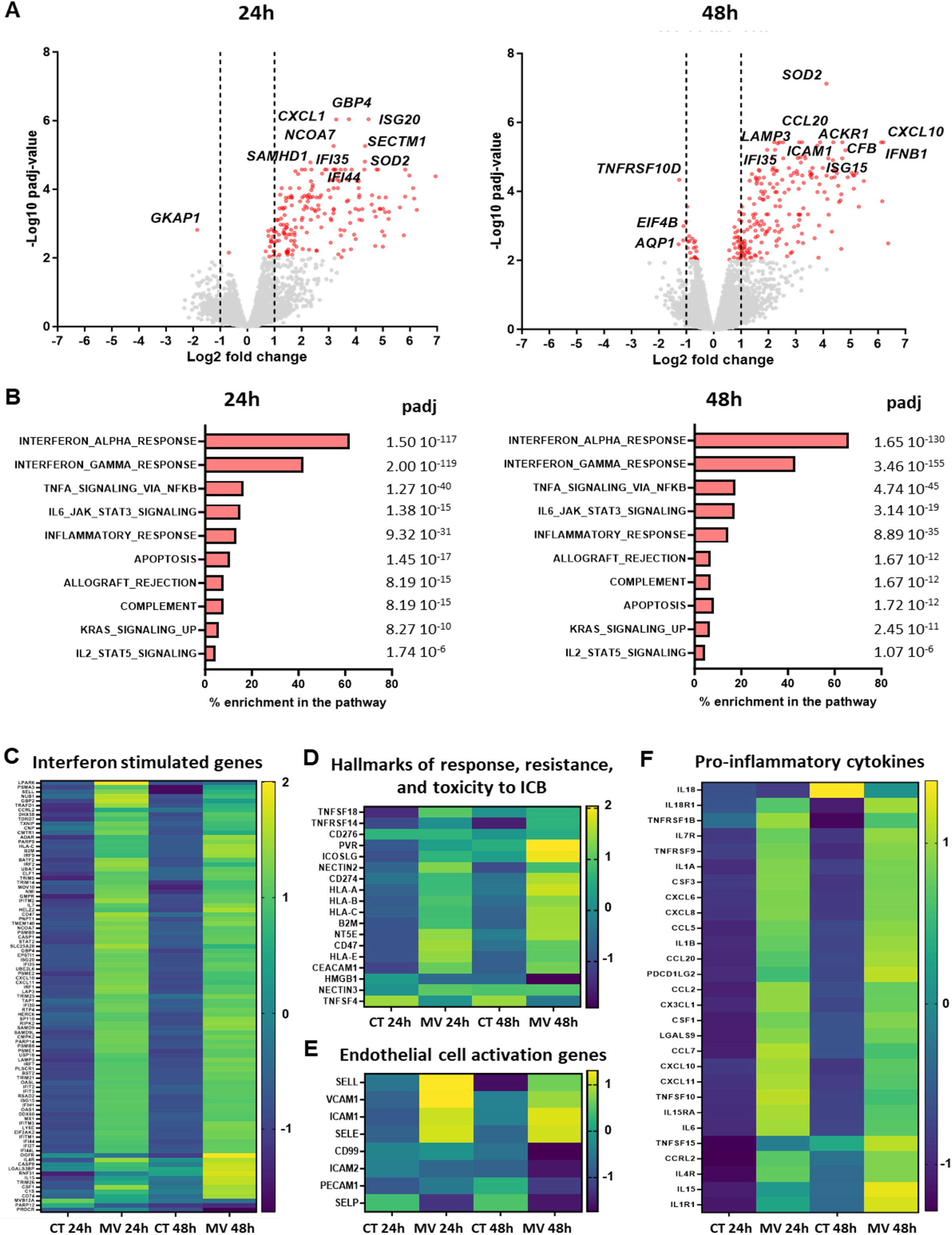
MV infection induces modification of the TME at the transcriptomic level. Chips were infected with 1x10^6^ MV particles on day 5 of the MVN formation by administration of the virus in the lateral media channel. mRNA of whole VMOCs were extracted at 24h and 48h post infection and analyzed using 3’ RNA-seq. A) Volcano plots of differentially expressed genes. Red: p≤ 0.01; Grey: p>0.01. Dotted lines correspond to fold change equal to 2 or -2. B) GSEA Pathway enrichment analysis. C-F) Heatmaps of the expression of interferon-stimulated genes (ISGs) (C), of hallmarks of response, resistance, and toxicity to immune checkpoint blockade (ICB) (D), of endothelial cell activation genes (E) and of genes encoding for pro-inflammatory cytokines (F).

## Discussion

Much work has focused on elucidating the mechanisms underlying tumor-selective replication of the vaccinal Schwarz strain of measles virus (MV) in pleural mesothelioma and other cancers. In contrast, the mechanisms governing viral interactions with the TME and the physical and biological barriers that influence viral activity remain less characterized. In this work, we studied MV oncolytic activity and its consequences on the TME through the intravascular delivery using our *in vitro* VMOC model.

We first optimized the experimental conditions to generate an endothelial compartment with two PVs connected to a central MVN. We showed the importance of the two-step method and of the endothelial cell/fibroblast ratio to obtain an optimal number of bridging vessels for an optimal circulation of molecules and particles through the vascular network. A deep characterization of this compartment demonstrated the presence of adherens and tight junctions between endothelial cells in both the PV and MVN, as well as the formation of a complex network with numerous branches in the MVN. PVs also exhibited functional responses, including modulation of permeability in response to Ang-1 and elongation under flow. The analysis of the cellular heterogeneity using scRNA-seq highlighted the presence of subpopulations of the different cell types used to constitute VMOC with similar signatures to those found in previous studies. Interestingly, we observed that the addition of mesothelioma cells in VMOC influenced the heterogeneity of cells in the TME, the main consequence being a decrease of the basal type I and II interferon pathway. Finally, administration of MV into the endothelial compartment led to the infection and death of PM cells. Infection of fibroblasts was also observed along with transcriptional evidence of induction of ISGs, activation of inflammatory signaling, endothelial activation, and changes in overall immunogenicity.

Fibroblasts are major components of the TME where they differentiate into CAFs. These cells contribute to tumor progression, resistance to therapies, and angiogenesis. Moreover, fibroblasts have been widely used *in vitro* to promote MVN formation and stability through their pro-angiogenic signaling (39,45,47,61). Fibroblasts have also been reported to support the formation of a normal vasculature (62). Here, we observed that the ratio of ECs to hLFs was critical for the formation of functional and perfusable MVNs. Low fibroblast density resulted in poorly organized networks with larger vessel diameters, whereas high fibroblast density led to the formation of smaller vessels with limited connectivity to PVs. This effect of lung fibroblasts on MVN architecture was already observed in a previous endothelial–fibroblast co-culture model, in which increased fibroblast density was directly proportional to reduced vessel diameter (26). To confirm the quality of the endothelial network, the permeability of the endothelial barrier to 70-kDa FITC-dextran was evaluated. The permeability coefficient (Pd) measured for PVs was consistent with what is described in literature (37,33). Unfortunately, the complexity of the MVN did not allow an accurate measurement of Pd; however, the presence of adherens and tight junctions, criteria already used in other studies (63,64), support the quality of the network.

CAFs exist as different subpopulations with pro- or anti-tumor properties (65). The scRNA-seq analysis revealed that different subpopulation of hLFs were present in VMOC with some of them displaying transcriptomic signatures close to previously described myCAF and particularly to ECM-myCAF populations (56). Addition of PM cells did not drastically impact fibroblasts heterogeneity; however, decreased expression of genes associated with the IFN-I response was observed, which could explain their increased susceptibility of hLFs to MV infection in the VMOC. In contrast, hLFs in 2D monoculture were minimally infected (Figure S15), consistent with our previous observations in CCD-19Lu human lung fibroblasts, where resistance to infection was associated with a functional IFN-I response (6). Reduced baseline expression of antiviral and IFN-I–related genes in CAFs has been reported and linked to tumor-derived TGF-β signaling (66). PM cells are known to produce TGF-β (67,68), and scRNA-seq CellChat analysis in our model identified all PM cell subpopulations as the predominant source of TGF-β signaling toward endothelial and fibroblast populations (Figure S9A-panel i), consistent with this hypothesis.

A similar reduction in the IFN-I transcriptomic signature was observed in endothelial cells upon co-culture with PM cells, accompanied by the disappearance of the EC_Interferon subpopulation. However, despite these transcriptomic changes, endothelial infection in the VMOC remained limited. An explanation could be that the proximity between PM cells and fibroblasts is higher in VMOC compared with ECs. Modifications to EC activation has already been observed in 2D co-culture with lung cancer cells (24,69,70) but not in the context of PM, in complex 3D models or at the level of the subpopulation. Furthermore, these consequences for ECs and fibroblasts may vary depending on the PM cell used, as it was already shown with lung cancer cells (24,70). Overall, these data highlight the importance of complex co-culture models for studying OV activity, as they enable the integration of intercellular signaling that influences viral susceptibility.

As expected, an induction of the expression of genes coding for IFN-I (*IFNB1*), and of a high number of ISGs was obtained after MV infection, including genes coding for chemotactic mediators (*CXCL10*, *CXCL11*), viral RNA sensors (*DDX58*, *DHX58*, *IFIH1*), translation inhibitors (*IFIT1*/*2*/*3*/*5*), RNA-degradation enzymes (*OAS1*/*2*/*3*, *OASL*), and restriction factors (*MX1*/*2*, *BST2*, *RSAD2*, *IFI6*/*27*). This ISG induction is consistent with our previous observations in monocultures of PM cells (10) and in co-culture systems with myeloid cells (17). The mechanisms are well-described and involve innate immune sensors such as retinoic acid-inducible gene I (RIG-I) and melanoma differentiation-associated gene 5 (MDA5) that detect viral RNA in cells (71). In addition, a strong induction of several inflammatory cytokines was observed after MV infection with notably increased levels of CXCL10 (IP-10), a chemokine involved in the recruitment of immune cells in the TME. Induction of IP-10 is frequently observed after virus infection and has also been reported in complex co-culture models (20,21). This increase in IP-10 is at least partly due to infected mesothelioma cells, as we have already demonstrated (17,72). An activation of endothelial cells was also observed, with the induction of the expression of genes coding for proteins involved in the adherence and rolling of circulating immune cells. Similar transcriptional signatures have been reported in an MVN-on-chip system exposed to SARS-CoV–2 (20). In addition, genes coding for proteins involved in antigen presentation were also induced in the VMOC. Altogether, these effects should contribute to the recruitment of immune cells and promote an anti-tumor immune response. In support of this, a recent vascularized skin-on-chip model showed that IL-8 induction following herpes simplex virus (HSV) infection drove neutrophil migration from the vascular channel toward infected keratinocytes, while IL-8 neutralization reduced vascular adhesion and transendothelial migration (73). Given that IL-8 is strongly induced after MV infection in the VMOC model, recruitment of neutrophils could be expected if they are introduced into our vascular compartment.

An increase in the expression of genes coding for proteins involved in the inhibition of immune responses, such as *CD274* (PD-L1), *CD47,* and *HLA-E*, were also observed. This suggests that combination with immune checkpoint inhibitors (ICIs) may improve the anti-tumor immune effects of MV. Anti-PD-1–blocking antibodies are relevant in this context, as they are already used as first-line therapy for PM in combination with anti-CTLA-4 (1). Furthermore, activation of the RIG-I and/ MDA5 pathways has been reported to enhance the efficiency of ICIs, including anti-PD-1 (74,75). RNA viruses such as MV naturally activate these pathways, providing a mechanistic rationale for combination strategies.

As mentioned above, though limited infection was observed, the endothelial compartment in the VMOC was affected by MV and led to the induction of genes coding for proteins involved in immune cell recruitment. However, a disruption of vessel architecture was also observed, in particular in PVs and with the condition 2x10^6^ viral particles (Figure S11D). Vascular network-on-chip models infected with H1N1, or SARS-CoV-2 have already shown structural disruption despite low levels of infection (20,21). In the latter study, the authors reported that paracrine IFN-I signaling from infected organoids co-cultured with the MVN mediated endothelial disruption without direct infection, which may explain our observation. Indeed, a strong activation of the type I IFN pathway was observed after MV infection in the VMOC, in addition to severe inflammation, which together could be comparable to a cytokine storm as proposed by the authors using H1N1 or SARS-CoV-2 viruses (20,21).

Recently, vascularized models of PM-on-chip were used to assess the response to CAR-T cells using patient derived mesothelioma samples (76). Building on this approach, the VMOC model could potentially be adapted at a translational level to evaluate therapeutic responses and combination strategies. In addition, VMOC could be useful to decipher resistance mechanisms to therapy and/or the contribution of the cells from the TME in the response to treatment, given its tunable cellular composition. The VMOC could also be interesting for the study of intercellular communication within the TME in a controlled and more integrated setting. This is supported by our observation that the incorporation of mesothelioma cells into the MVN modified cellular heterogeneity and transcriptional profiles.

In conclusion, the VMOC model provides a platform to study the response to oncolytic viruses across multiple components of the TME, including tumor, vascular, and stromal compartments. The consequences of virus infection in the model are consistent with known antiviral and inflammatory responses to MV and other viruses. This model may therefore serve as a useful platform for investigating resistance mechanisms and evaluating novel therapeutic combinations, beyond oncolytic viruses, to improve antitumor efficacy.

## Acknowledgements

The authors thank the IBISA MicroPICell facility (Biogenouest) and the member of the national infrastructure France-Bioimaging supported by the French National Research Agency (ANR-10-INBS-04) for technical support, for their expert technical assistance. We are most grateful to the Genomics and Bioinformatics Core Facility of Nantes (GenoBiRD, Biogenouest) for its technical support.

## Author contribution

NR, JF, MW, and HP performed experiments; OH and BVS analyzed vessel permeability; ED, MD, and NR performed scRNASeq experiments, LT performed scRNAseq analysis, LT and IC provided a strong expertise on endothelial cell to analyze and interpret the data; NB and JFF provided a strong expertise on oncolytic viruses to design experiments, analyze and interpret the data; NR, DF, NB, JFF, LT, BVS and CB interpreted the study and provided critical review; CB and BVS conceived, designed, supervised, analyzed, interpreted the study. NR, BVS and CB wrote the manuscript. All authors reviewed the manuscript.

## Funding

This work was supported by INSERM, CNRS, the French national research agency (ANR, project n° ANR-21-CE06-0034-04) and Ligue Régionale contre le cancer Grand-Ouest.

## Notes

### Competing Interest Statement

The authors have declared no competing interest.

