## Supplementary figures for "Oncolytic measles virus reprograms the tumor microenvironment in a vascularized mesothelioma-on-chip model"

#### Supplementary figure legends

**Supplementary figure S1.** Characterization of the permeability and response to stimuli of the parental vessels (PV). A) Permeability measurement of PV. (i) Representative fluorescence image of 70 kDa FITC–dextran diffusion in the chip. Donor voxel in PV and receiver voxel in gel (white dashed squares) indicate the regions used for fluorescence intensity quantification. Voxel size = 50 × 50 × 250 $\mu$m. The area between voxels (orange line) defines the diffusion path across the barrier. Paired donor–receiver voxels were selected at three different diffusion sites located between the lateral and central channels per chip. (ii) Equation used to calculate the permeability coefficient (P, cm/s), as previously described by Tu *et al.* (31). Fixed parameters for the equation were: t1 (time point 1) = 60 s, t2 (time point 2) = 300 s, $V_{receiver}$ (reciver voxel volume) = 6.25x10-7 $cm^3$, $A_{barrier}$ (barrier area between donor and receiver) = $1.25x10^{-4}$ $cm^2$, $V_{receiver}/A_{barrier}$ = 0.005 cm. $I^{t1}_{donor}$, $I^{t2}_{donor}$, $I^{t1}_{receiver}$ and $I^{t2}_{receiver}$ represent fluorescence intensities measured in donor and receiver voxels at $t_1$ and $t_2$. (iii) Permeability coefficient (P, cm/s) measured under three conditions: absence of PV (no HUVECs), presence of PV (with HUVEC) and PV with Angiopoetin 1 (Ang1) treatment (HUVEC+Ang1). Bars represent mean ± standard deviation. n = 2 (no HUVECs), n = 3 (with HUVECs), and n = 1 (HUVECs + Ang1) chip. B) Response of PVs to the application of a flow of medium for 24h using the 4U Microfluidic Pressure Pump (Cellix, cat. no. 4U-EC-PC). Upper panels: Brightfield microscopy images; lower panel: Confocal microscopy images. Green, VE-cadherin; purple, nucleus.

**Supplementary figure S2.** Optimization of the microvascular network (MVN) by one-step method. A) Schematic of MVN formation by one-step method in the idenTX chip. Grey, MVN; purple; Fibrin gel. B) Representative confocal microscopy images showing MVN obtained using 8, 10, 12 and $15x10^6$ HUVECs/mL (upper panels) and after 70 KDa FITC-dextran injection into the PVs. Grey, HUVECs mRuby; Green, 70 kDa FITC dextran. Scale bar = 200 $\mu$m. C) Graph showing average lumen size of the microvessels in the MVN obtained with various HUVECs quantity. Horizontal bars represent mean ± SEM (n=2 chips, 3 ROIs (Region of interest) of 1272×1272 $\mu$m per chip). D) Lumen size distribution of microvessels in the MVN. Graph represents mean ± SEM (n=2, 3 ROIs of 1272x1272 $\mu$m for each chip). E) Percentage of open bridging vessels obtained with different quantities of HUVECs loaded in the central gel channel. Bars represent mean ± SEM. (n=2 chips, 1 ROI of 1767 × 8134 $\mu$m per chip).

**Supplementary figure S3.** Methods to measure open bridging vessels and size of lumen of the microvascular network (MVN) in the CC. A) Schematic of VMOC showing bridging vessels connecting PV and MVN. B) Representative confocal microscopy image showing the difference between open and closed bridging vessel. Grey, HUVECs; Green, 70 kDa FITC dextran. Scale bar = 200 $\mu$m C) Image processing for lumen size measurement, as previously described by Wan *et al.* (24). (i) A representative ROI (1272 × 1272 $\mu$m) from one chip was used for lumen size quantification. (ii) Lumen diameters in the MVN were measured using the ImageJ distance measurement tool. Three ROIs were analyzed per chip, and the mean value was calculated for each chip. Scale bar = 100 $\mu$m.

**Supplementary figure S4.** Two-step method for generating a perfusable microvascular network (MVN) connected to parental vessels (PV). A) Schematic of the two-step method. The experimental procedure consists of injecting a first layer of HUVECs (15 × $10^6$ cells/mL) to promote the formation of bridging vessels, followed by injection of a second layer of HUVECs (10 × $10^6$ cells/mL) with primary human lung fibroblasts (hLFs; 1 × $10^6$ cells) to induce branched MVN formation. B) Representative confocal microscopy images showing MVNs generated using the two-step method (left panels) and after 70 kDa FITC-dextran injection into PVs (right panels) on day 5 post seeding. Grey, HUVECs mRuby; Green, FITC dextran 70 kDa. Scale bar = 200 $\mu$m. C) Graph showing the average lumen size of microvessels within the MVN obtained using the two-step method with varying hLF numbers. Horizontal bars represent mean ± SEM. n=3 chips, 3 ROIs of 1272x1272 $\mu$m for each chip. **$p<0.01$; *$p<0.05$. Unpaired t-test (two-tailed). D) Lumen size distribution of microvessels within the MVN obtained using the two-step method. Graphic represents mean ± SEM. (n=3 chips, 3 ROIs of 1272x1272 $\mu$m for each chip).

**Supplementary figure S5.** Characterization of the microvascular network (MVN) obtained using two-step method. A) Microscopy images showing the evolution of MVN formation over 10 days. Scale bar = 500 $\mu$m. B) Representative confocal microscopy images of the MVN showing VE-cadherin, CD31, and ZO-1 immunostaining on day 5.

**Supplementary figure S6.** Transcriptional heterogeneity and functional characterization of pleural mesothelioma (PM) cells in the VMOC model. A) UMAP representation and distribution of all cells in the VMOC with (Meso, n=2) or without (No_Meso, n=2) PM cells. B) t-SNE projection of PM cells subtypes across two biological replicates (Meso_1 and Meso_2). C) Relative proportion of Stress, Proliferative, and OXPHOS cancer cell subtypes across the two biological replicates. D-F) Gene set enrichment analyses across PM cell transcriptional states (Proliferative, OXPHOS, and Stress), with each state compared against all other cancer cell states.

**Supplementary figure S7.** Transcriptional heterogeneity and functional characterization of endothelial cells in the VMOC model. A) UMAP projection of endothelial cells (EC) showing distinct transcriptional states, including EC_Interferon, EC_Artery, EC_Inflam, EC_Mesenchymal, and EC_Proliferative, in four VMOC, including with (Meso, n=2) and without PM (No_Meso, n=2). B) Relative proportion of endothelial cell states across VMOCs with and without PM, highlighting shifts in endothelial composition in the presence or absence of PM cells. C-D) Gene set enrichment analysis in the two endothelial cell states (EC_Proliferative and EC_Mesenchymal) under No_Meso conditions, with each EC cluster compared against all other EC clusters. E) Feature plots of endothelial cell population with marker genes for proliferative (*MKI67*, *TOP2A*) and mesenchymal endothelial cell states (*PTX3*, *ESM1*). F) Reactome pathway enrichment analysis of the EC_Interferon endothelial cell state, with pathways enriched relative to all other endothelial cell clusters. G) Heatmap of interferon-stimulated gene expression across all endothelial cell states. H) Gene set enrichment analysis of all endothelial cells in the Meso condition relative to No_Meso.

**Supplementary figure S8.** Fibroblast heterogeneity and trajectory analysis in the VMOC model. A) UMAP projection of fibroblast showing distinct transcriptional states, in four VMOC, including with (Meso, n=2) and without (No_Meso, n=2) PM cells. B) Relative proportion of fibroblasts cell states across VMOCs with PM cells (and without PM cells. C) Left: Integrated UMAP of cells (endothelial cells and fibroblasts) colored by annotated cell states. Right: Pseudotime analysis showing inferred cellular progression across the integrated cell populations. D) Gene set enrichment analysis of all fibroblast populations in the Meso condition relative to No_Meso.

**Supplementary figure S9.** Cell–cell communication networks of TGFβ and NOTCH signaling pathways inferred from scRNA-seq. A) TGFβ signaling network. (i) Heatmap of cell–cell communication probabilities in the TGFβ signaling network, with cell subpopulation as senders (rows) and receivers (columns). (ii) Bar plot showing the relative contribution of individual ligand–receptor pairs to the overall TGFβ signaling network. Contributions are calculated as the ratio of the total communication probability of each ligand–receptor pair to the total communication probability of the TGFβ signaling pathway. B) NOTCH signaling network. (i–ii) Same as in (A), for the NOTCH signaling pathway.

**Supplementary figure S10.** Evaluation of MV infection after administration in the central gel channel of the VMOC on day of gel seeding. A) Representative confocal microscopy images of infection on day 5 after administration of MV directly in the central gel channel at the time of gel loading on day 0 and at different multiplicity of infection (MOI). B) Representative confocal microscopy images of infection on day 5. Meso13 cells were pre-infected in 2D for 2h at MOI 1 prior to incorporation into the central gel channel during gel loading on day 0. Green: GFP expressing cells (infected cells); red: Meso13; yellow: infected Meso13; grey: HUVECs mRuby. GFP signal detected in regions lacking overlap with both PM (red) and HUVEC (grey) markers was considered indicative of infected fibroblasts. Scale Bar = 500 $\mu$m.

**Supplementary figure S11:** Optimization of MV-GFP infection parameters (dose and exposure time) via the lateral media channel in the VMOC. A) Representative confocal image showing MVN collapse when medium is not refreshed for more than 24 h. Chips were maintained under static conditions from day 5 to day 8 of MVN formation. B) Representative confocal images of MV-GFP infection at increasing viral doses following 24 h exposure. Chips were exposed to $2 \times 10^5$, $5 \times 10^5$, or $1 \times 10^6$ $TCID_{50}$ of MV-GFP, followed by medium replacement. Images were acquired at 48 h post infection. Scale bar: 200 $\mu$m. C) Representative confocal image at 24 h post infection following exposure to $1 \times 10^6$ $TCID_{50}$ of MV-GFP for 6 h. D) Representative confocal image at 48 h post infection following exposure to $2 \times 10^6$ $TCID_{50}$ of MV-GFP for 6 h. Scale bar: 200 $\mu$m. Green: GFP expressing cells (infected cells); red: Meso13; yellow: infected Meso13; grey: HUVECs mRuby. GFP signal detected in regions lacking overlap with both PM (red) and HUVEC (grey) markers was considered indicative of infected fibroblasts.

**Supplementary figure S12.** Cytokine concentrations measured in culture supernatants of VMOC 48h after MV infection. Cytokine concentrations were measured using LEGENDplex assay. Graphics represent mean ± SEM of 3 independent experiments. *, $p<0.05$; ***, $p<0.001$. Unpaired t test (two-tailed).

**Supplementary figure S13.** Transcriptional effects of MV in VMOC. A) Venn diagram representing the number of genes commonly and specifically deregulated 24h and 48h after MV infection. B) Pathway enrichment analysis of genes commonly deregulated 24h and 48h after MV infection.

**Supplementary figure S14.** Proportions of endothelial and fibroblast cell states estimated from bulk 3′RNA-Seq data following MV infection using single-cell–derived reference signatures. A) Relative proportions of endothelial cell states estimated across conditions (Ctrl and MV-infected Endo and Meso conditions at 24h and 48h). Endo refers to cells from the parental vessels, whereas Meso corresponds to cells from the central channel. The left panel shows mean proportions, while the right panel shows mean ± SEM. B) Relative proportions of fibroblast states estimated across conditions (Ctrl and MV-infected Endo and Meso at 24h and 48h). C) GSVA scores of endothelial cell state signatures across conditions (Ctrl and MV-infected Endo and Meso at 24h and 48h). D) GSVA scores of fibroblast state signatures across conditions.

**Supplementary figure S15.** Infection of hLFs, HUVECs, and Meso13 cells in monoculture by measles virus (MV). Cells were seeded at 10,000 per well in 24-well plates and infected the following day with MV-GFP (MOI = 1). Top (brightfield) and bottom (GFP fluorescence) panels show representative images acquired at 72 h post infection.

### Supplementary Figure S1

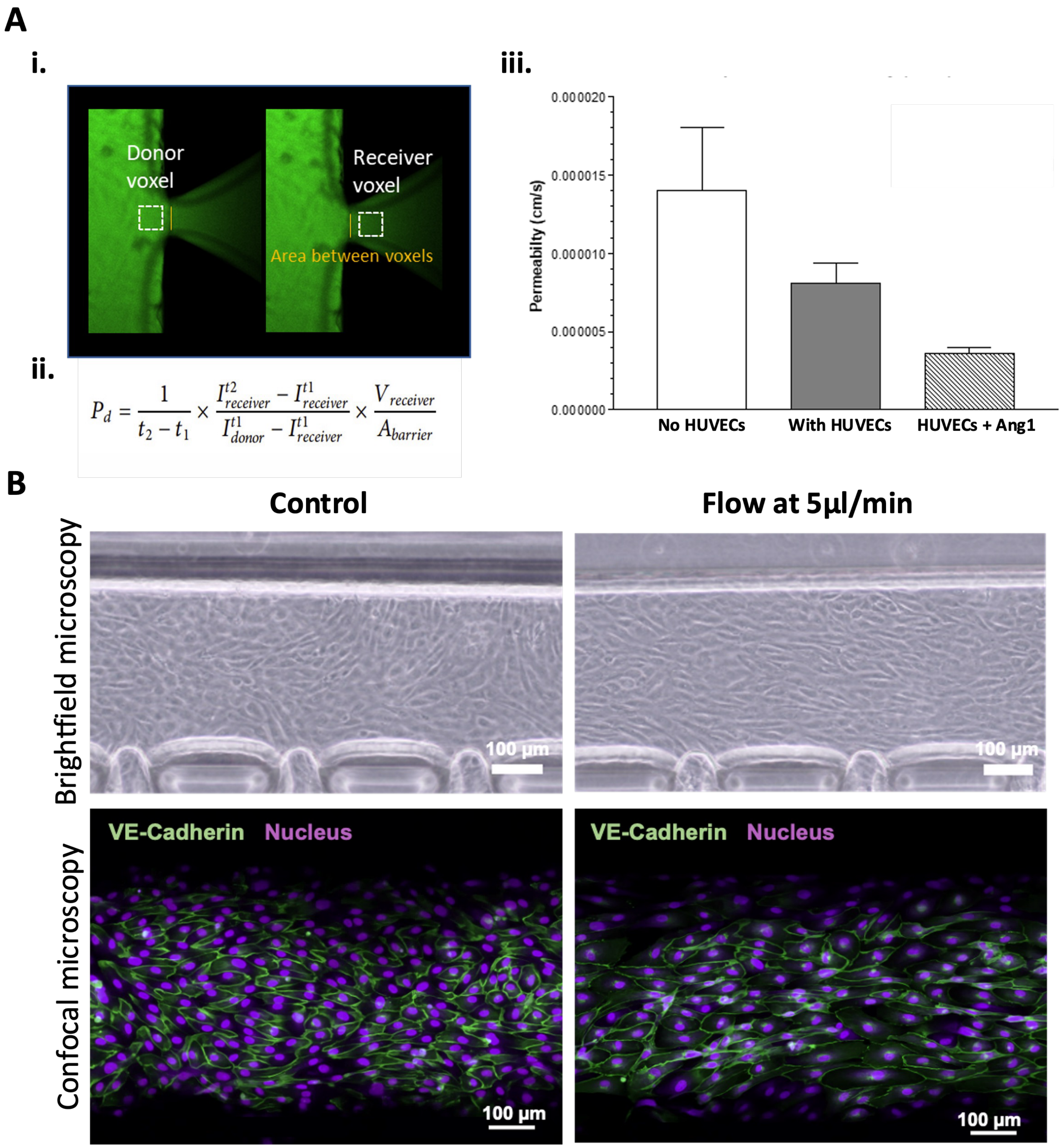

#### Supplementary Figure S2

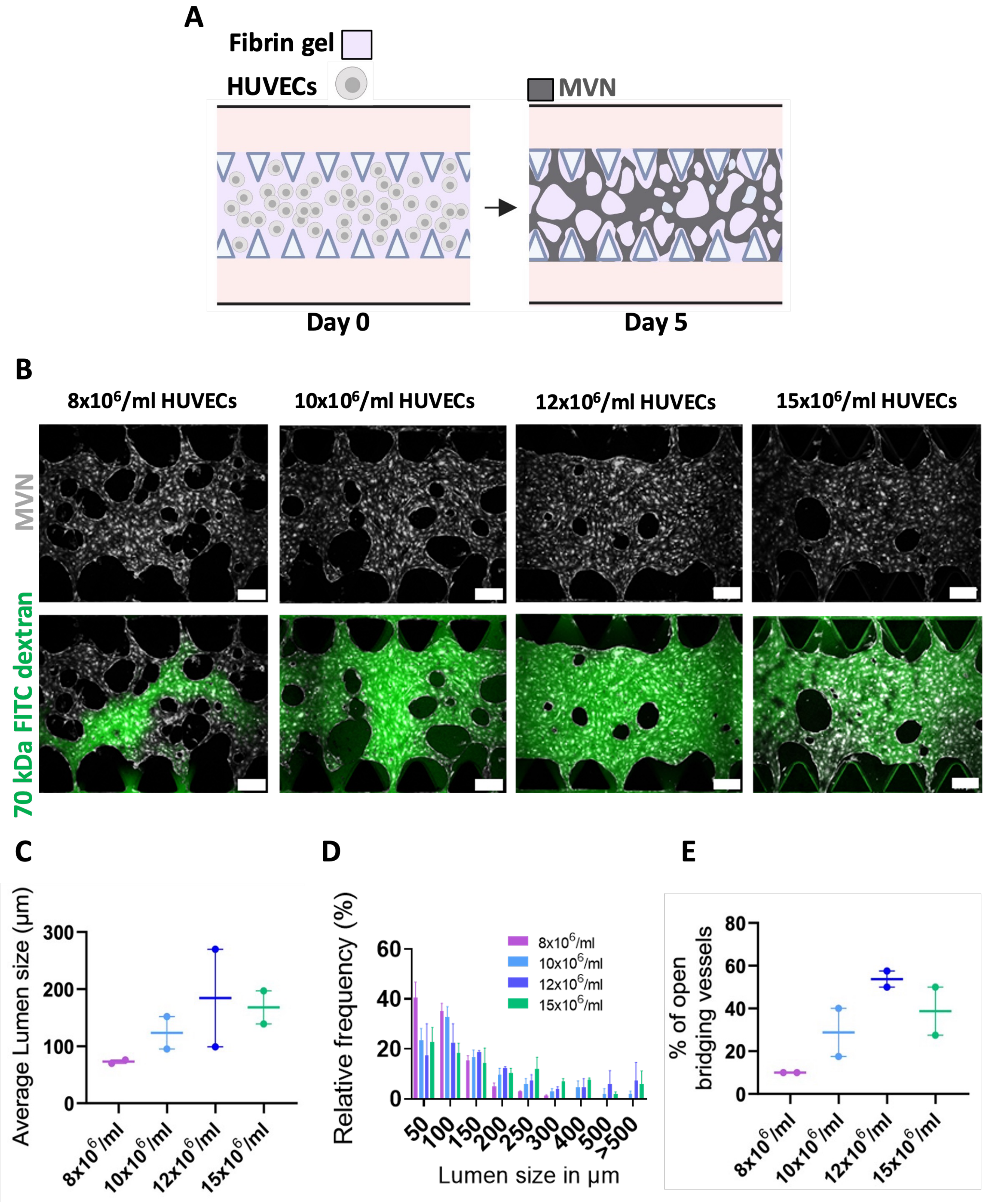

**Supplementary Figure S3**

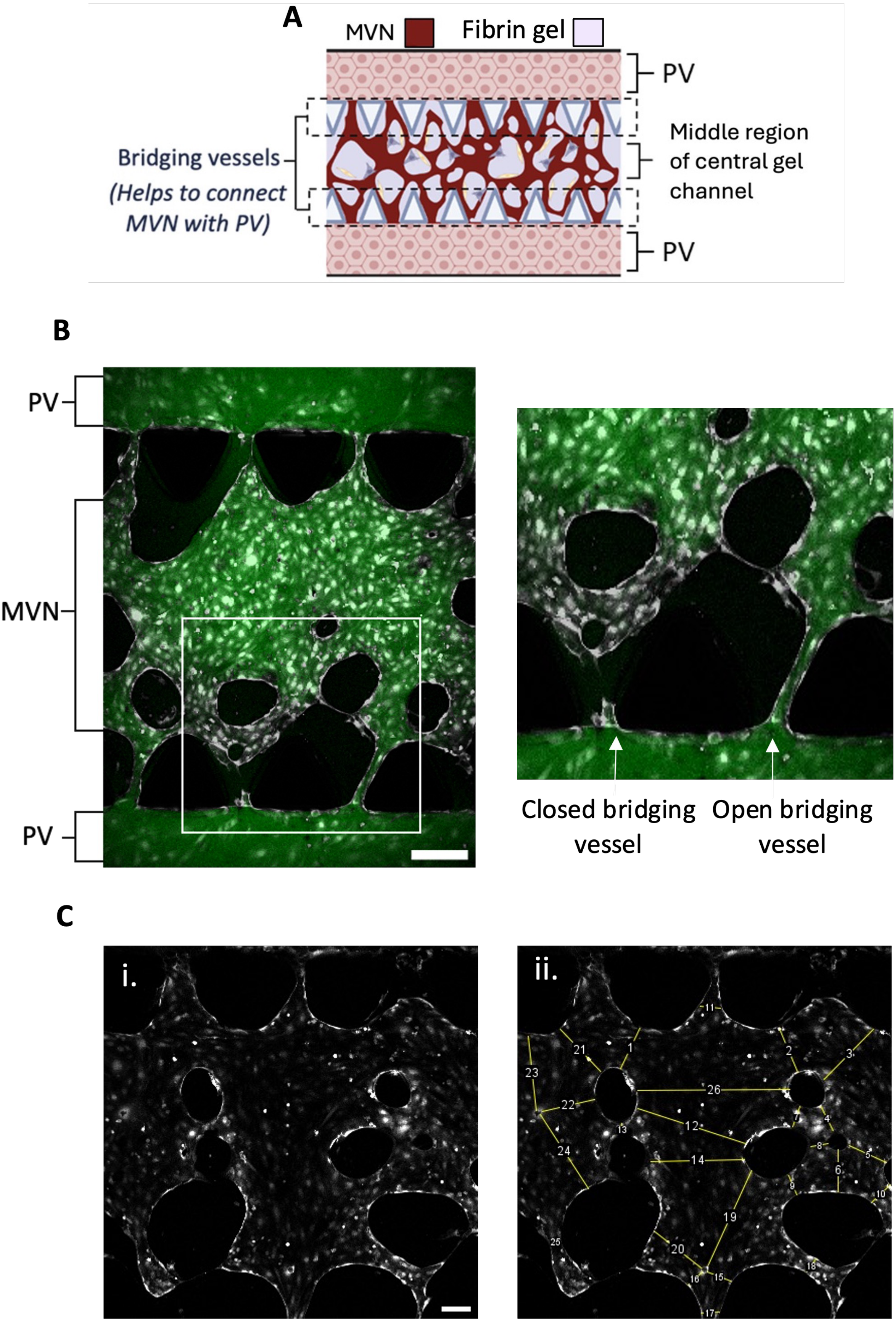

#### Supplementary Figure S4

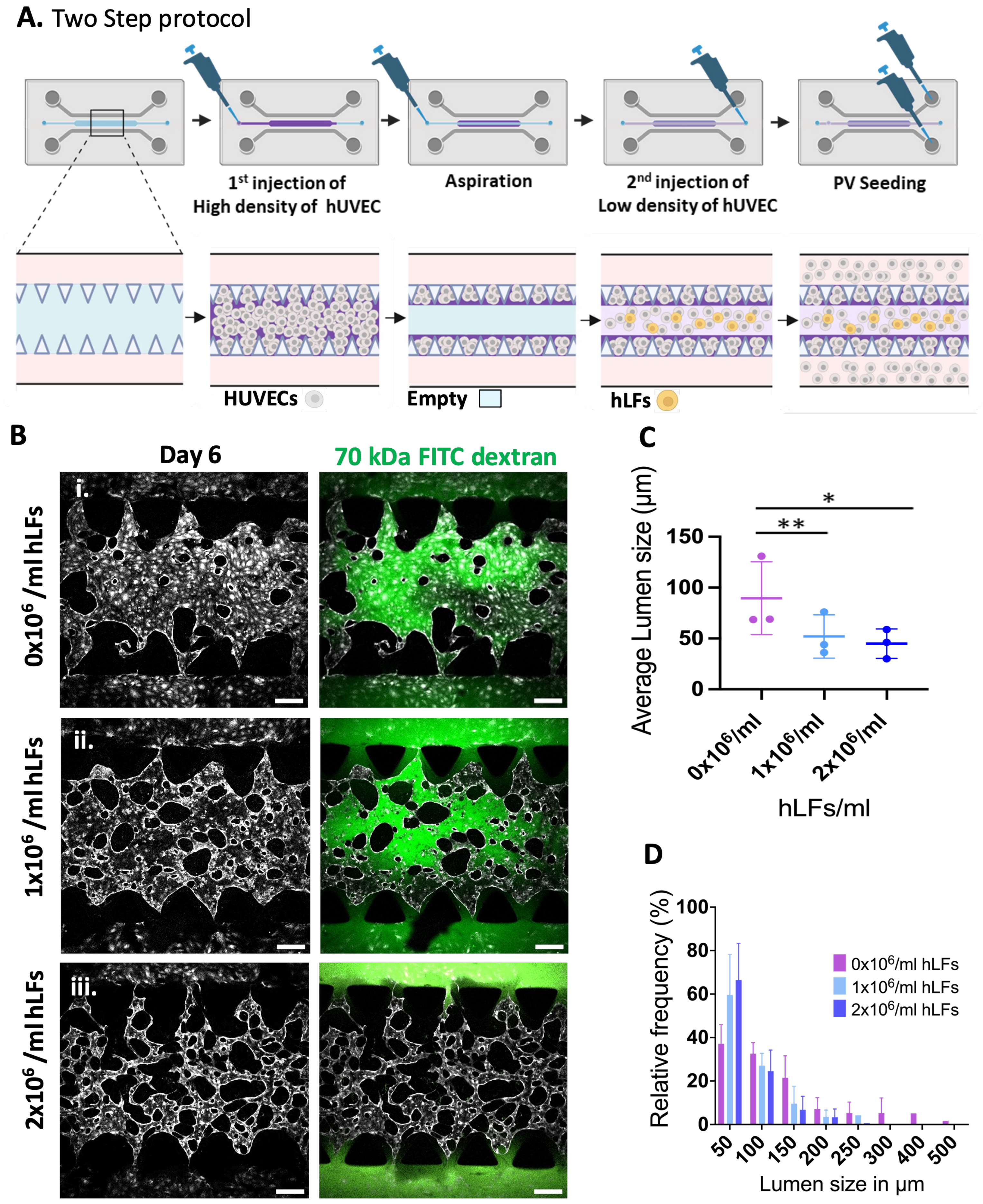

### Supplementary Figure S5

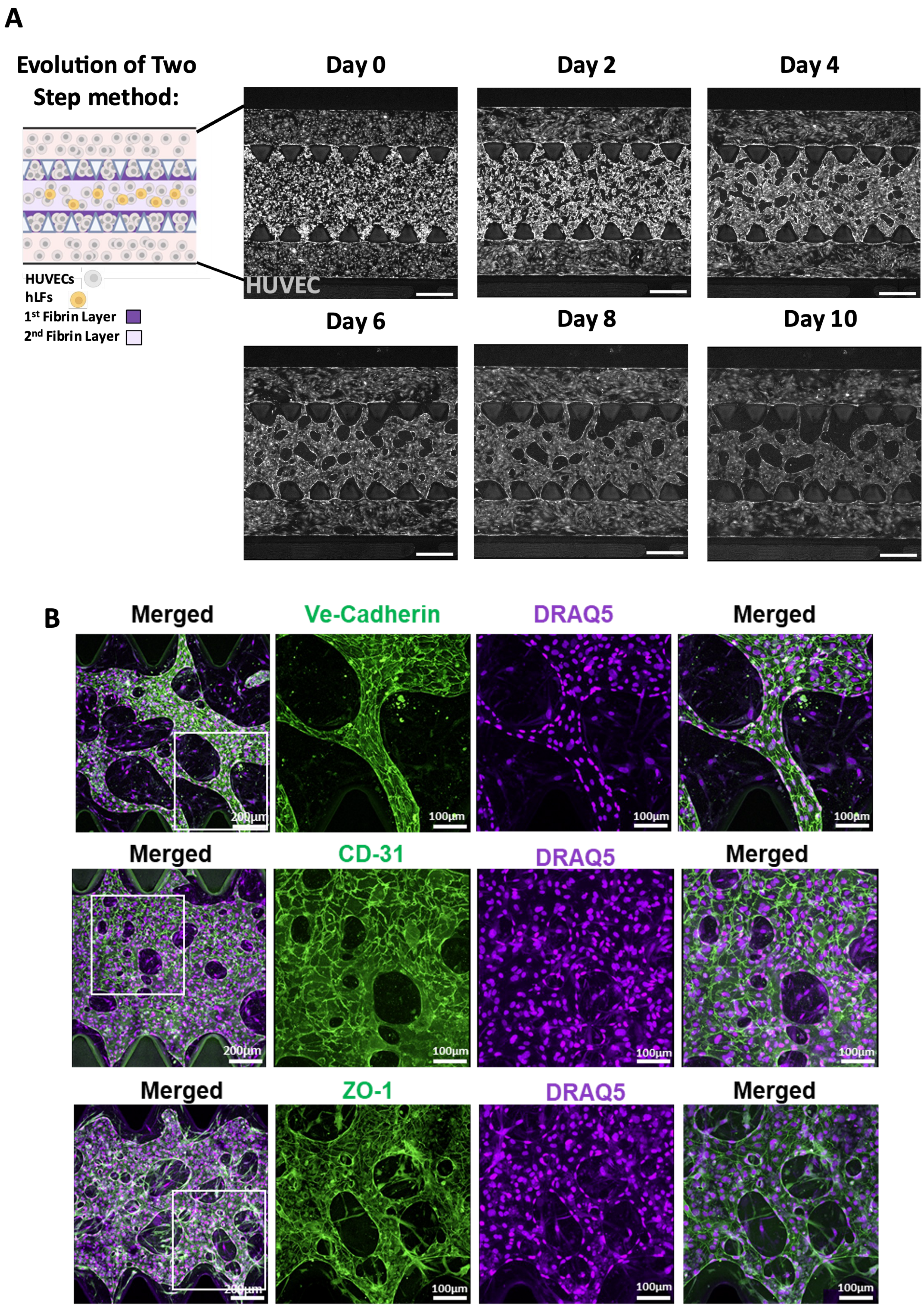

### Supplementary Figure S6

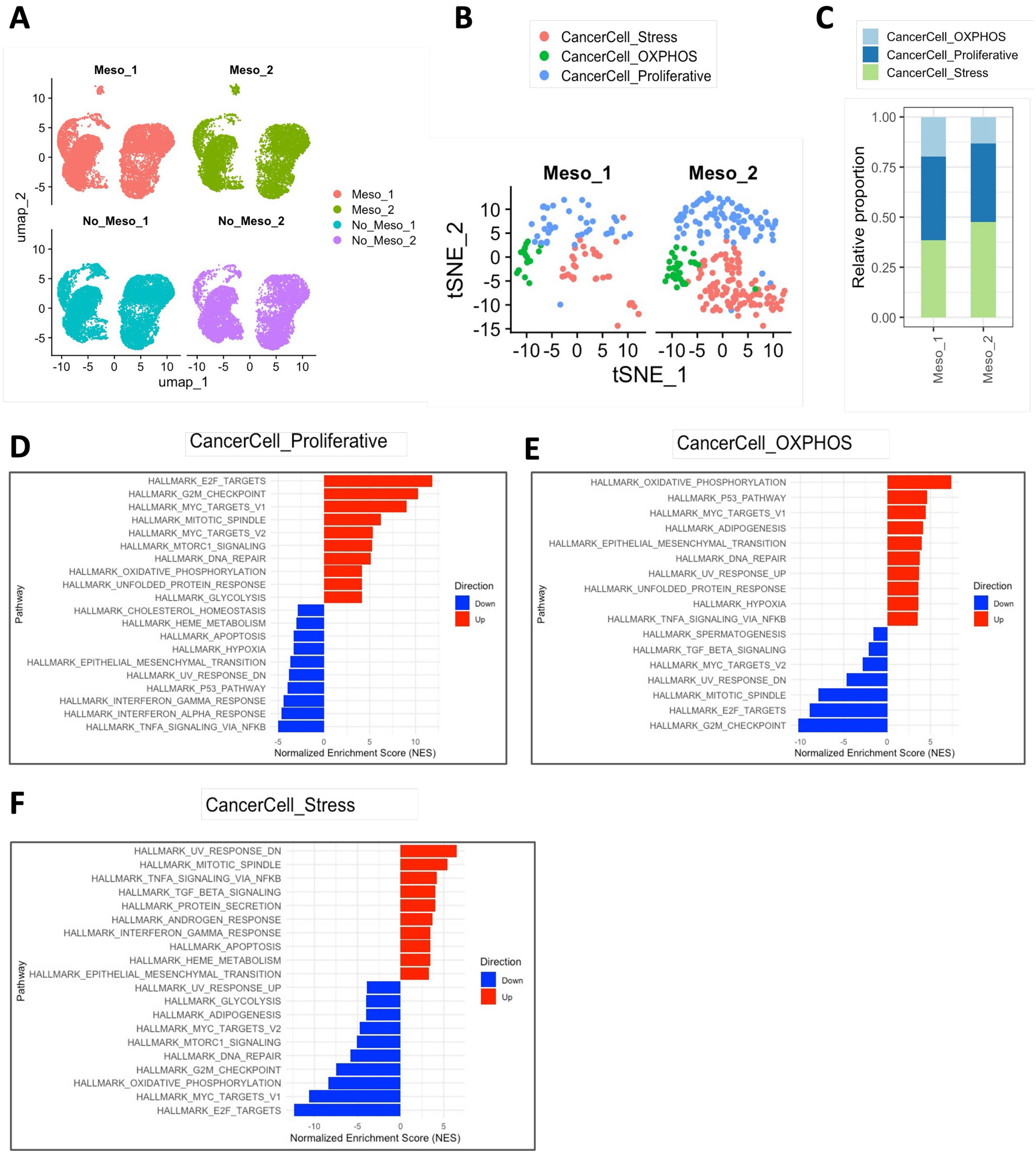

### Supplementary Figure S7

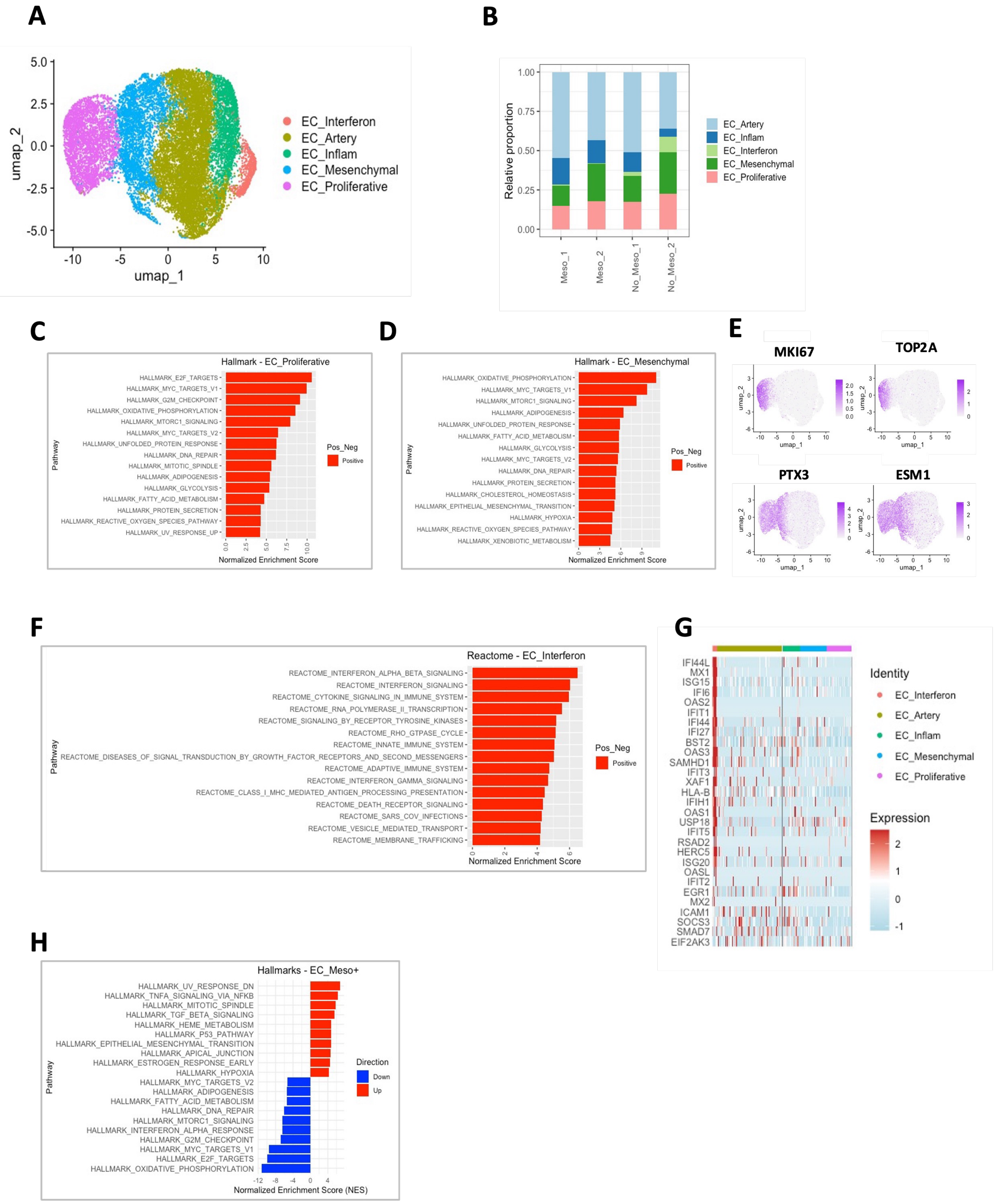

#### Supplementary Figure S8

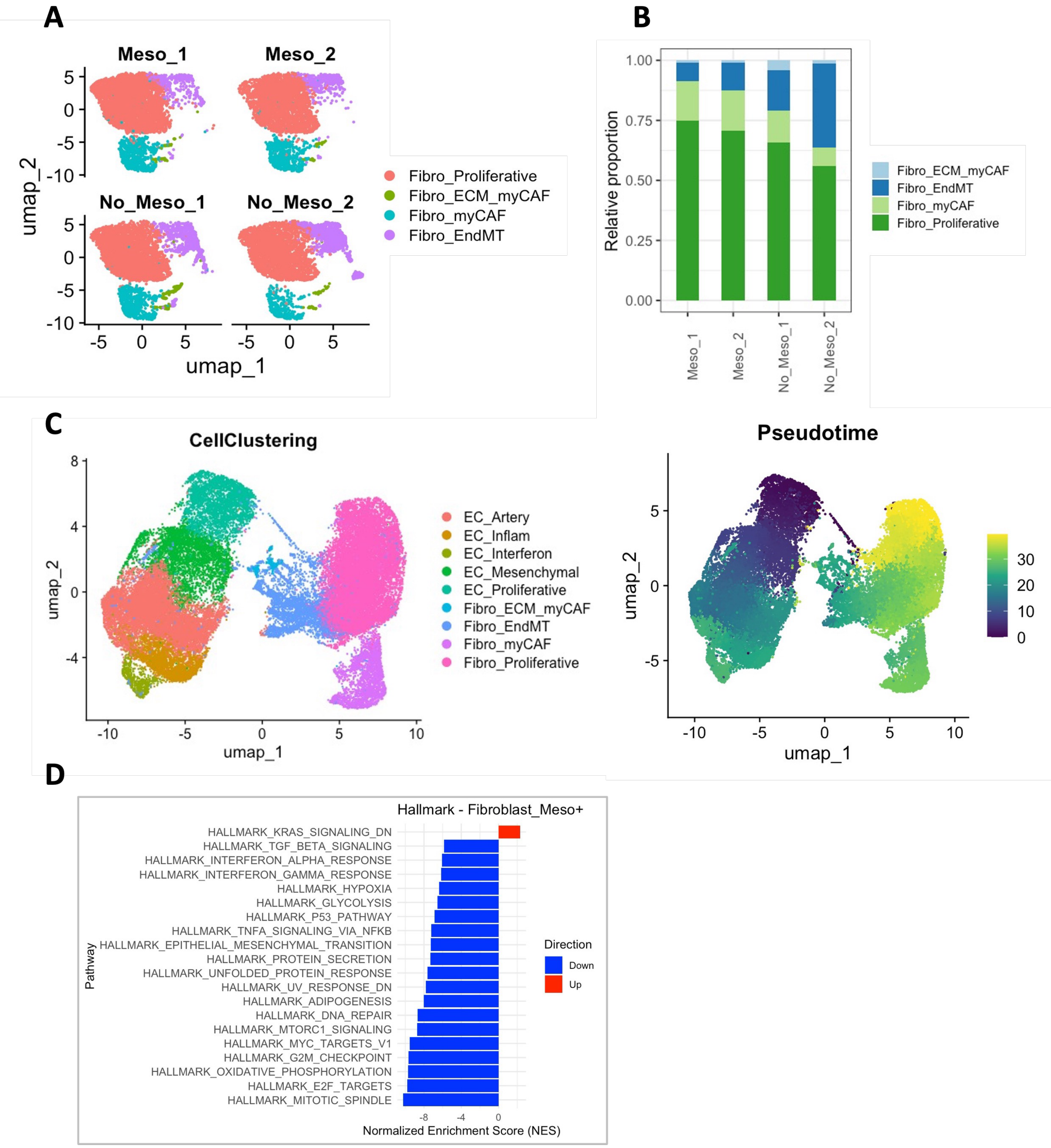

### Supplementary Figure S9

## A

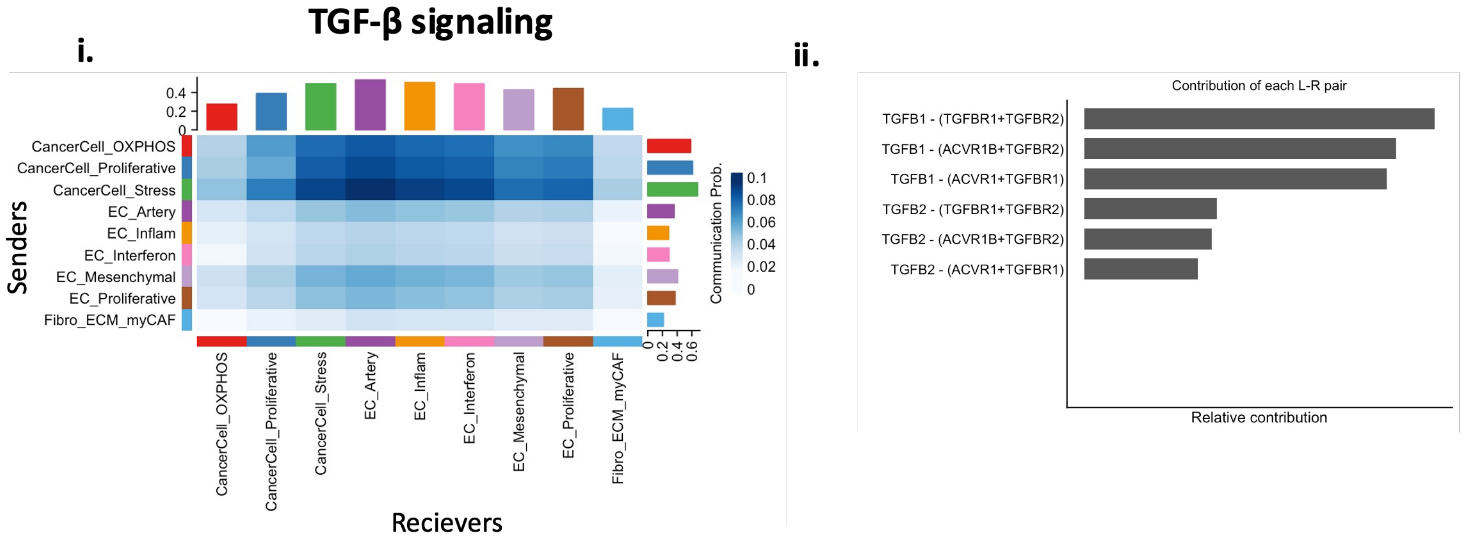


## B

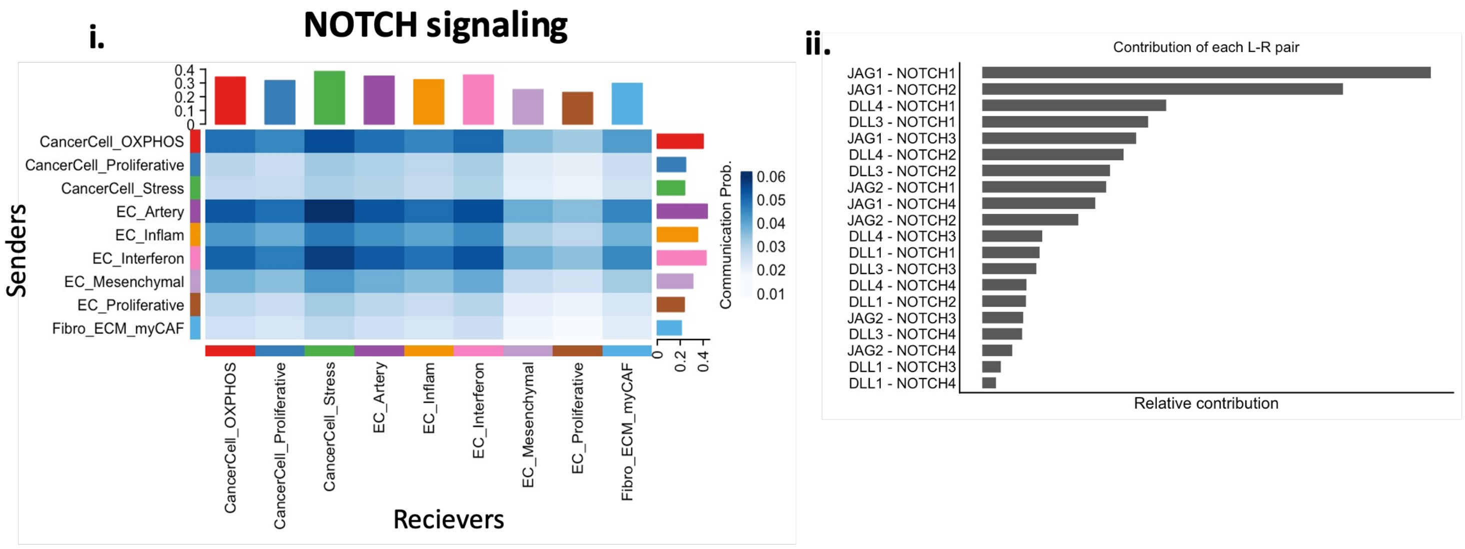

### Supplementary Figure S10

**A**

**MOCK infection** **MOI 1** **MOI 10**

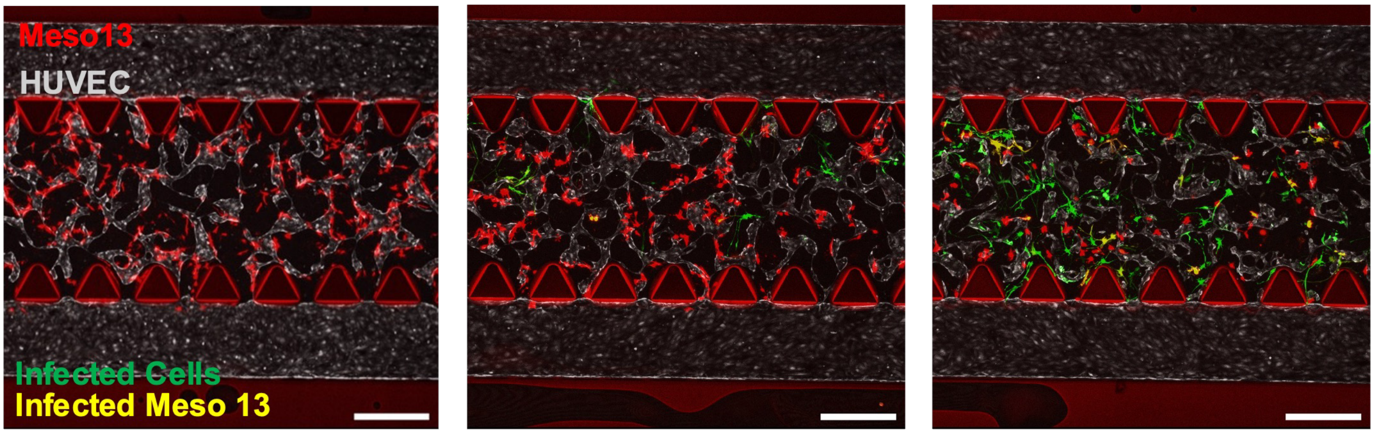


**B**

**Preincubation with MOI 1**

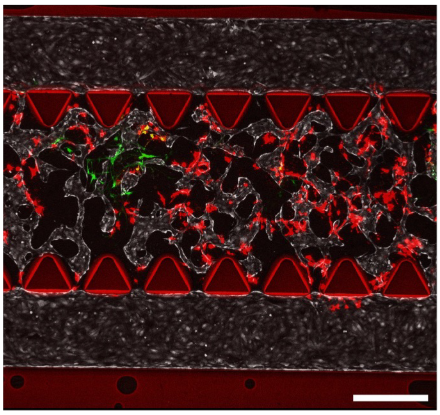

### Supplementary Figure S11

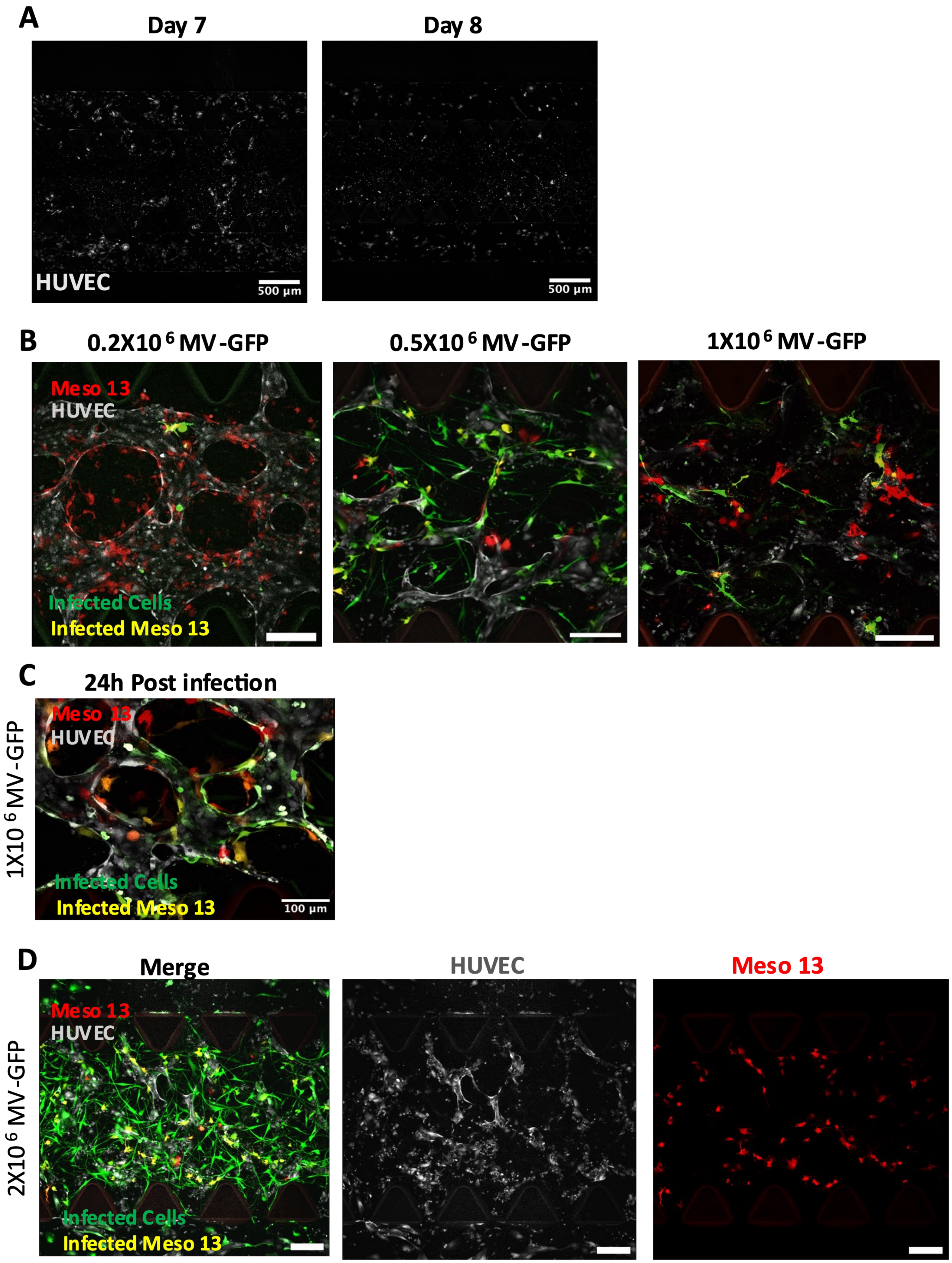

### Supplementary Figure S12

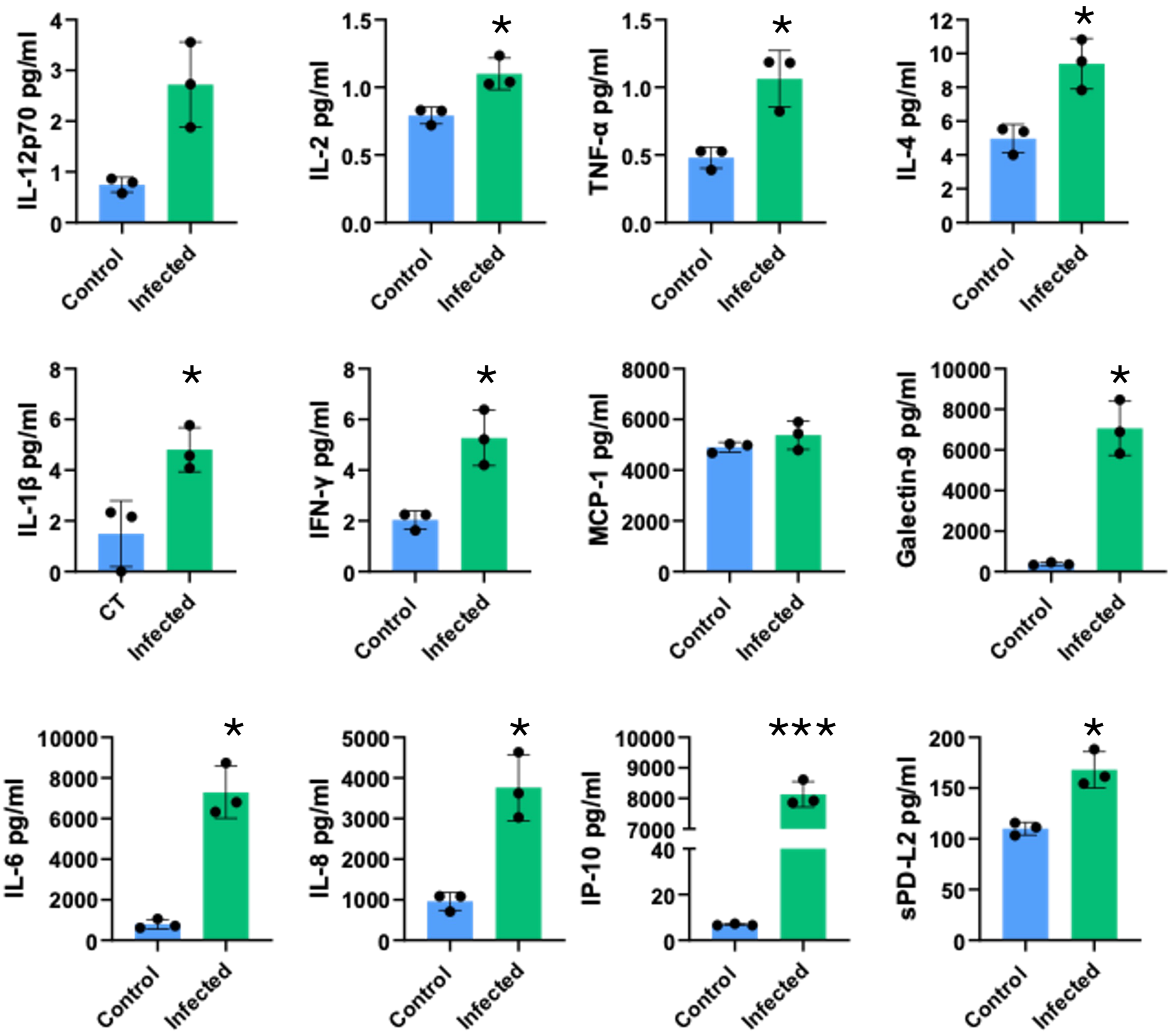

#### Supplementary Figure S13

A

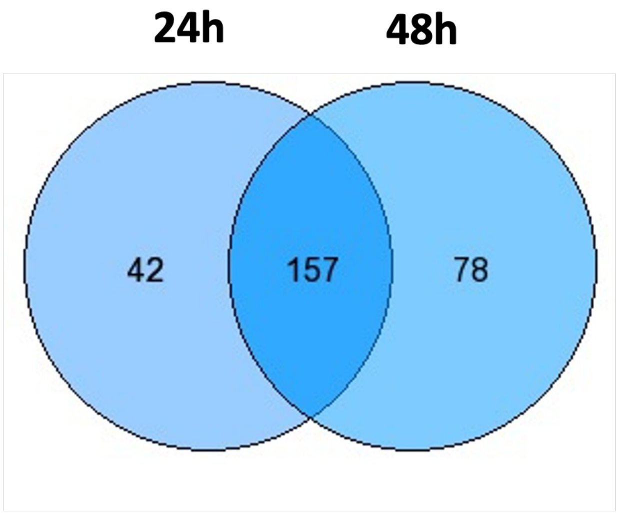


B

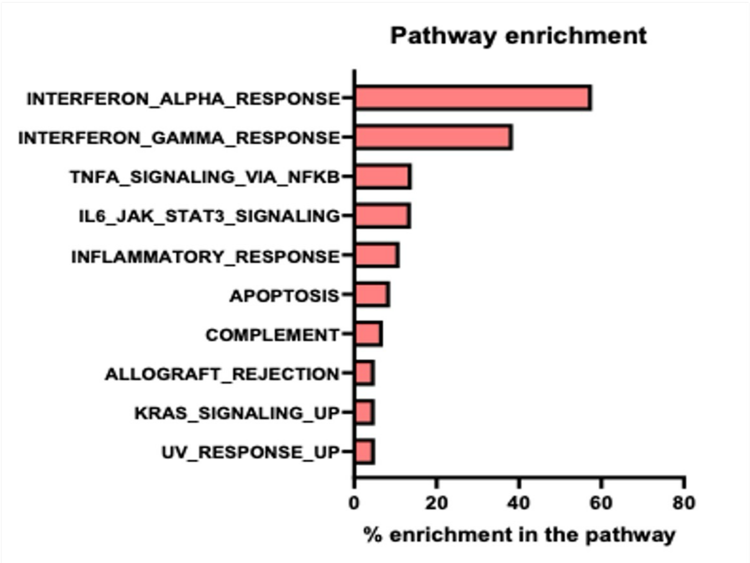

### Supplementary Figure S14

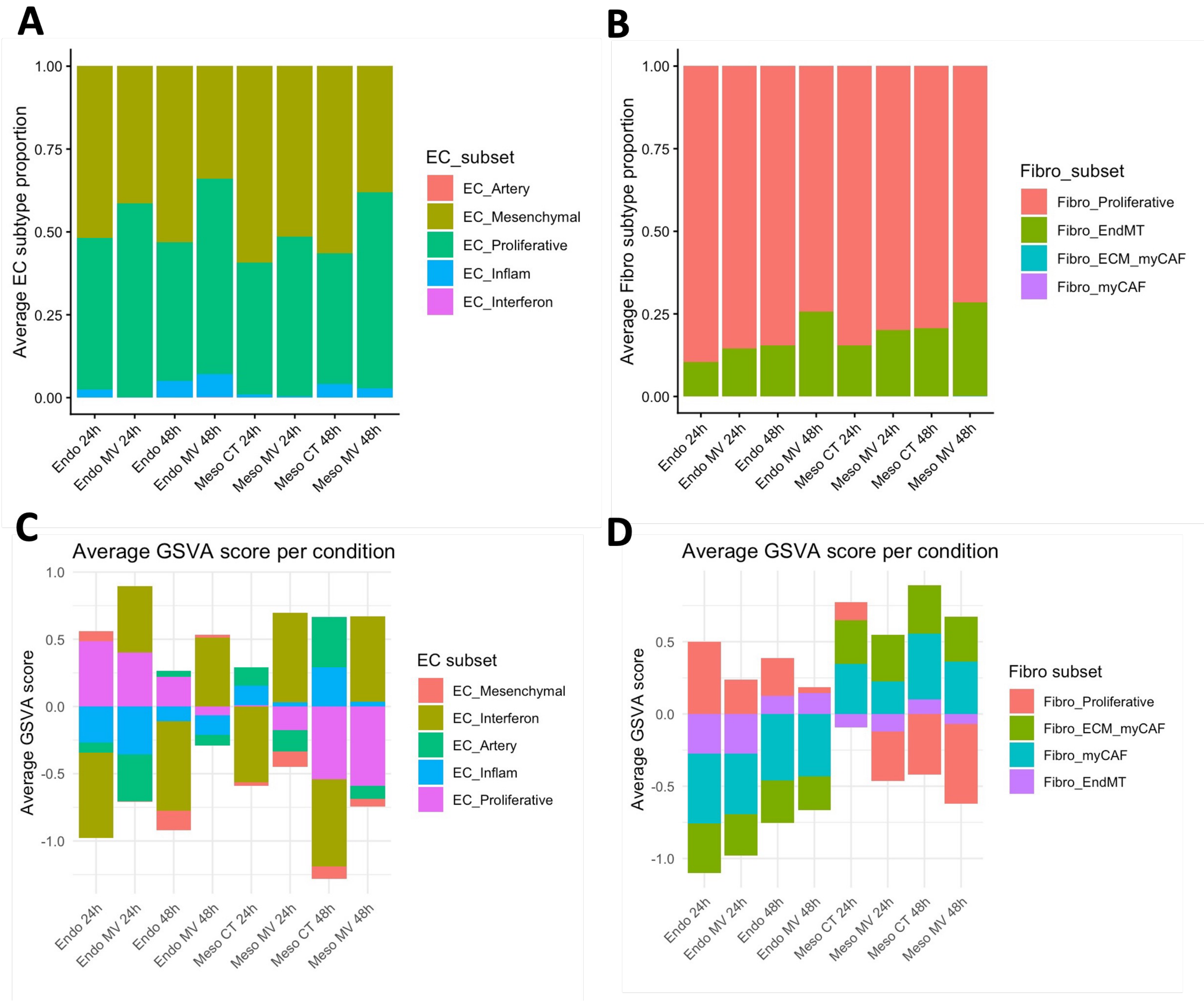

### Supplementary Figure S15

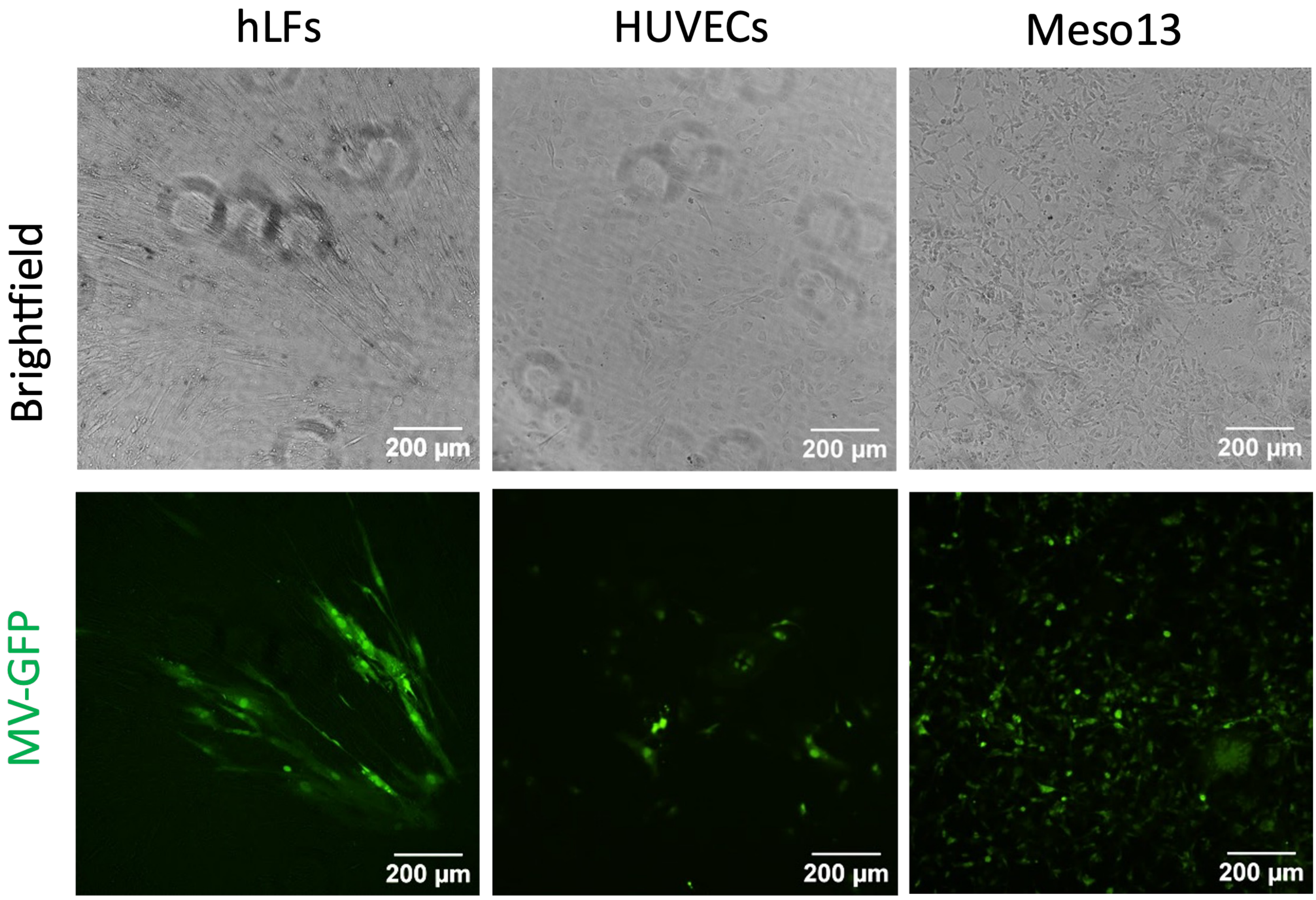
