## Supplementary figures and images for "Oncolytic measles virus reprograms the tumor microenvironment in a vascularized mesothelioma-on-chip model"

### Supplementary movie S1

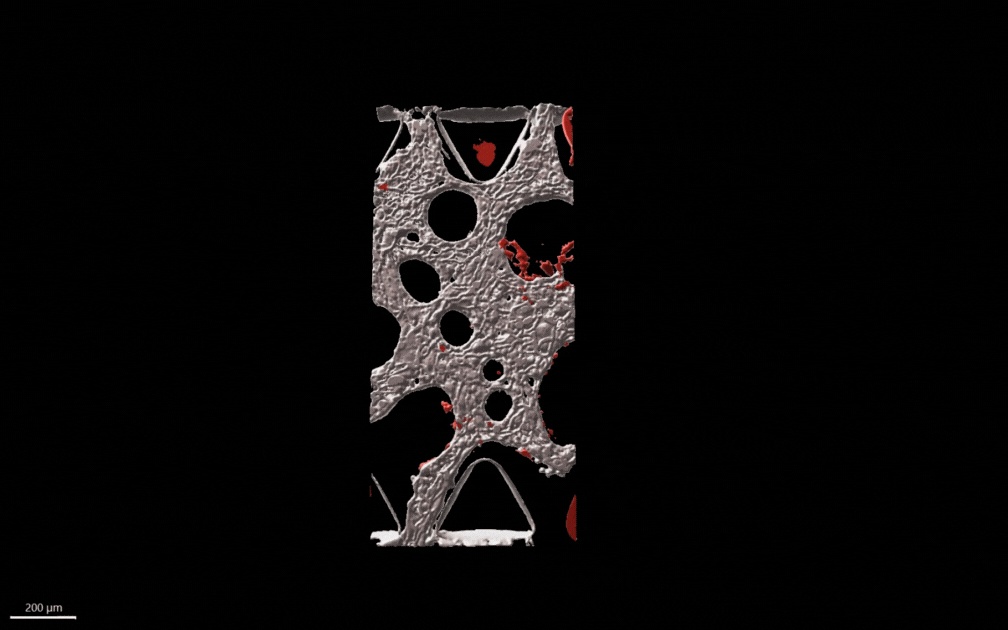

### Supplementary movie S2

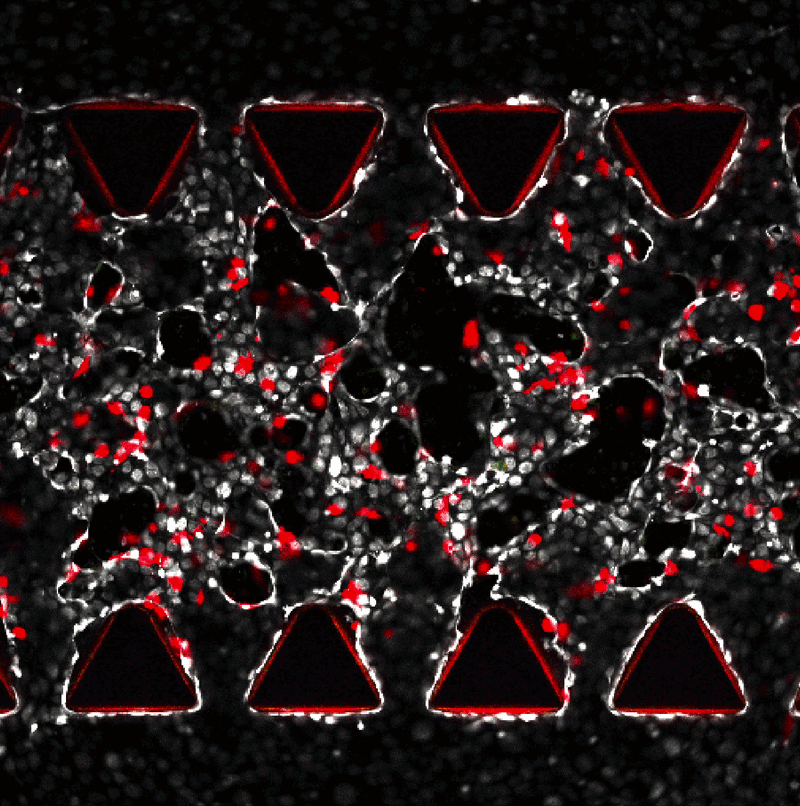
